# A Streamlined High-Throughput LC-MS Assay for Quantifying Peptide Degradation in Cell Culture

**DOI:** 10.1101/2024.10.11.617883

**Authors:** Samuel J. Rozans, Abolfazl S. Moghaddam, E. Thomas Pashuck

## Abstract

Peptides are widely used in biomaterials due to their easy of synthesis, ability to signal cells, and modify the properties of biomaterials. A key benefit of using peptides is that they are natural substrates for cell-secreted enzymes, which creates the possibility of utilizing cell-secreted enzymes for tuning cell-material interactions. However, these enzymes can also induce unwanted degradation of bioactive peptides in biomaterials, or in peptide therapies. Liquid chromatography-mass spectrometry (LC-MS) is a widely used, powerful methodology that can separate complex mixtures of molecules and quantify numerous analytes within a single run. There are several challenges in using LC-MS for the multiplexed quantification of cell-induced peptide degradation, including the need for non-degradable internal standards and the identification of optimal sample storage conditions. Another problem is that cell culture media and biological samples typically contain both proteins and lipids that can accumulate on chromatography columns and degrade their performance. However, removing these constituents can be expensive, time consuming, and increases sample variability. Here we show that directly injecting samples onto the LC-MS without any purification enables rapid and accurate quantification of peptide concentration, and that hundreds of LC-MS runs can be done on a single column without a significantly diminish the ability to quantify the degradation of peptide libraries. We also show that column failure is evident when hydrophilic peptides fail to be retained on the column, and this can be easily identified using standard peptide mixtures for column benchmarking. In total, this work introduces a simple and effective method for simultaneously quantifying the degradation of dozens of peptides in cell culture. By providing a streamlined and cost-effective method for the direct quantification of peptide degradation in complex biological samples, this work enables more efficient assessment of peptide stability and functionality, facilitating the development of advanced biomaterials and peptide-based therapies.

## 1 INTRODUCTION

Peptides are short protein sequences that can signal cells and are also natural substrates for cell-secreted enzymes.^1^ Proteases are the most common class of enzyme that degrades peptides by hydrolyzing the amide bonds between amino acids.^2^ Peptides can be easily synthesized and modified with a range of bioconjugate chemistries,^3^ and they are often incorporated into biomaterial systems to improve bioactivity and enable cell-mediated scaffold degradation.^4,5^ For instance, cells are unable to adhere to many of the polymers used in synthetic cell culture systems, and these polymers are typically functionalized with RGD peptides to increase cell attachment.^6^ Cells are also frequently cultured within hydrogels, which are highly hydrated networks of crosslinked polymers that are designed to mimic the extracellular matrix that surrounds cells within tissues.^7^ Crosslinking these polymer networks with peptides that can be cleaved by proteases enables cells to actively degrade their local matrix, which enables them to spread and migrate within the hydrogels.^8,9^ Proteolytic degradation of peptides has also been used to modulate the signaling environment around cells,^10^ and release growth factors tethered to the polymer matrix.^11^

There are over 600 human proteases,^12^ including the matrix metalloproteinases (MMPs) family of proteases which are commonly harnessed to induce degradation of biomaterial scaffolds.^13,14^ There are also numerous exopeptidases which can non-specifically degrade peptides that are incorporated into biomaterials or are used therapeutically.^15,16^ The most common method for quantifying peptide degradation by proteases involves synthesizing peptides with a fluorophore and quencher at each end, then incubating the conjugate with the protease(s) of interest. However, many proteases, including MMPs, have promiscuous substrate specificity and a single peptide sequence is often cleaved by many proteases.^17^ The total protease activity near a cell or within a tissue is also incredibly complex, as cells secrete dozens of proteases, each with their own range of substrate specificities, in addition to protease inhibitors, and other cofactors that can modulate protease activity.^18^ Exopeptidases target the terminal amino acids of peptides, and their kinetics are highly sensitive to the chemistry of the termini.^16^ This poses a challenge for quantification methodologies that require the ends of the peptide to be modified, such as fluorophore-quencher pairs.^19^ Peptides can be rapidly made on automated synthesizers, which enables the facile creation of peptide libraries to rapidly screen numerous sequences.^20^ However, quantifying degradation using fluorescence-based methods limits the number of analytes in a single sample due to the need for minimally overlapping fluorescence spectra.

Liquid chromatography-mass spectrometry (LC-MS) is a robust analytical tool that integrates liquid chromatography to separate out and mass spectrometry to facilitate the multiplexed quantification of peptide degradation within complex samples.^21^ In liquid chromatography (LC), a mixture is introduced to a chromatography column, where the analytes are separated based on a solvent gradient. This is then sent to a mass spectrometer (MS), which can quantify up to thousands of different peptides within a single run.^22^ Since the peptides are identified by their molecular weight, there is no need to label the molecules with fluorophores or other chemistries which may change degradation kinetic. Furthermore, a single LC-MS run can take less than ten minutes, which enables a high sample throughput.

A key challenge in using LC-MS to quantify the functional degradation of peptides during cell culture or within biological samples is that these types of media contain different classes of molecules that are detrimental to either liquid chromatography or mass spectrometry. Specifically, the presence of high concentrations of proteins and lipids will readily foul HPLC columns,^23^ and the non-volatile salts are detrimental reproducible ionization in mass spectrometry and accumulate on the mass spectrometer.^24^ A variety of methods exist to isolate the analyte(s) of interest from these complex mixtures;^25,26^ however they are often expensive and time consuming. Furthermore, these purification techniques typically need to be optimized for a single analyte, and studies utilizing a library of peptides with a range of physiochemical properties are challenging for these methods.

In this work we have optimized methods for quantifying the stability and degradation of peptide libraries within cell culture media (Figure 1A). Directly injecting cell culture media into an LCMS has several advantages over methods that utilize purification steps because it requires minimal sample preparation and greatly reduces the chances of sample loss prior to analysis (Figure 1B). To ensure reproducible results, we tested a series of internal standard candidates and identified sequences which were both minimally degraded and were well retained on the column. We removed the salts present in the samples by sending the first section of the chromatography run to waste, which bypasses the mass spectrometer. A downside to directly injecting cell culture media onto the column is that the presence of lipids and proteins within the cell culture media degrades the performance of chromatography columns. Here we show that a single LC column can give reliable results for hundreds of runs without any sample purification and we identify the characteristics that are hallmarks of column degradation (Figure 1C). This method has benefits to existing purification protocols, such as solid phase extraction, in that it is less expensive, takes less time, and minimizes sample loss, even when studying complex peptide libraries. LCMS is a widely available technique, and these methodologies can be easily adopted by any laboratory with access to LCMS. Overall, this will help to advance our quantitative understanding of how cells degrade peptides, and improve our ability to design peptides that improve the efficacy of biomaterials and biomedical therapies.

**Figure 1.**
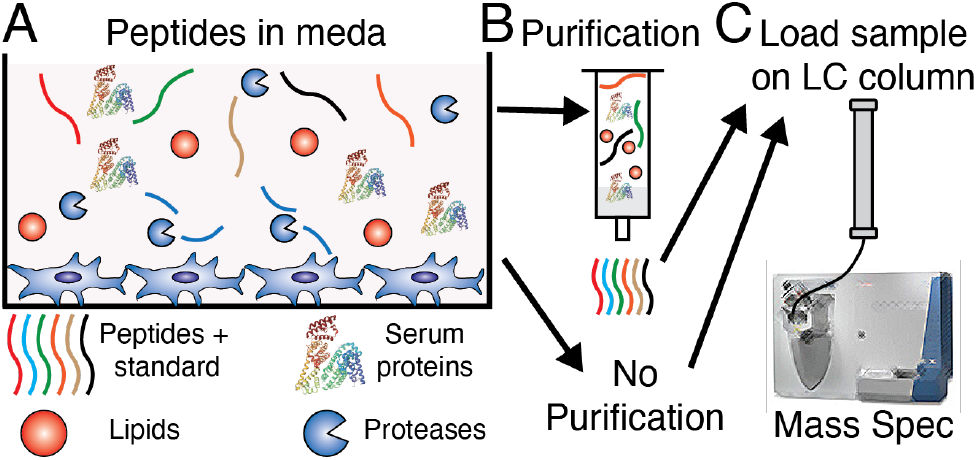
(A) Peptides and internal standards are added to cell culture media that contains numerous proteins and lipids. (B) This picture can either be purified or remain unpurified prior to (C) running the sample on LC-MS, in which the mixture is loaded onto a column follow by mass spectrometry.

## 2 MATERIALS AND METHODS

### 2.1 Materials

All peptide synthesis reagents were purchased from Chemscene or Ambeed. N,N-dimethylformamide (DMF) and dichloromethane (DCM) (both from VWR BDH Chemicals), piperidine (Millipore Sigma), trifluoroacetic acid (TFA) (Millipore Sigma), diethyl ether (Fisher Scientific), N, N-Diisopropylethylamine (DIPEA) (VWR), were used as purchased.

### 2.2 Peptide synthesis procedure

Peptides were synthesized using standard solid phase peptide synthesis (SPPS) protocols standard Fmoc-protected amino acids (Chemscene) on a Rink amide resin (Supra Sciences). All amide couplings were done using O-(6-chlorobenzotriazol-1-yl)-N,N,N′,N′-tetramethyluronium hexafluorophosphate (HCTU) in DMF. For each coupling the amino acid, HCTU, and DIPEA were added in a 4:4:6 molar ratio to the peptide. During peptide synthesis a ninhydrin test (Anaspec) was performed after every addition to test for the presence of free amines. Upon a positive test, the coupling was replicated until the test was negative. After successful coupling, the Fmoc group was removed by washing the resin with 20% piperidine in DMF twice for 5 min. A ninhydrin test was performed to check for a positive result.

For split-and-pool steps, the resin was washed 3× with DMF and then the entire amount of resin was weighed on a scale. This number was divided by 22 and this was split 19 ways into 19 separate tubes to ensure equal splitting of the resin. All natural amino acids (A, D, E, F, G, H, I, K, L, M, N, P, Q, R, S, T, V, W, and Y) were coupled in 19 separate 15 mL tubes. Upon successful coupling of all 19 amino acids, the 19 unique peptides were re-combined to form a library. Once a peptide library was completed, it was capped with acetic anhydride (Sigma-Aldrich) in a 10:5:100 acetic anhydride:DIPEA:DMF solution twice for 5 min, and then a ninhydrin test was performed to check for complete capping of the free amines. After capping the resin was washed 3× with DMF and 3× with DCM.

Peptides libraries and all peptides containing tryptophan were cleaved using 92.5% trifluoroacetic acid (TFA), 2.5% H_2_O, 2.5% triisopropylsilane (TIPS), and 2.5% dithiothreitol (DTT). Peptides not containing a tryptophan were cleaved using 95% trifluoroacetic acid (TFA), 2.5% H_2_O, and 2.5% triisopropylsilane (TIPS). Peptides were typically cleaved for 2-3 hours at room temperature using approximately 25 mL of cleavage solution per mM of peptide. After cleaving peptide(s) from the resin, they were precipitated in diethyl ether, centrifuged for 5 minutes at 4,000 rpm, and the supernatant was discarded. The peptide pellet was washed with diethyl ether, centrifuged two more times, then dried. Once dry the peptide(s) were dissolved in water and neutralized with ammonium hydroxide prior to purification.

All peptides were purified using high performance liquid chromatography (HPLC) using a Phenomenex Gemini 5 μm NX-C18 110 Å LC Column 150 × 21.2 mm. Gradients were run from 95% Mobile Phase A (water with 0.1% TFA) and 5% Mobile Phase B (acetonitrile with 0.1% TFA) to 100% Mobile Phase B. A typical HPLC run featured a two-minute equilibration step, followed by a 10-minute ramp from 95% Mobile Phase A to 100% Mobile Phase B, and then two minutes of equilibration at 100% Mobile Phase B, before ramping back down to the starting conditions. Notably, the split-and-pool libraries were ramped up to 100% Mobile Phase B over two minutes, since these libraries consisted of approximately 19 different peptides which weren’t intended to be separated from each other. After purification all peptides were lyophilized and were ready to use.

### 2.3. Quantification of sample storage conditions on peptide degradation

Human mesenchymal stem cells (hMSCs) (Rooster Bio, LOT 310368) at passage 3 were seeded into 24 well plates at a seeding density of 75,000 cells per well in 1mL of RoosterBasal-MSC-CC (RoosterBio, SU-022) containing RoosterBooster-MSC (RoosterBio SU-003). After 24 hour the media was changed, and peptide library AcβA-RGEFV-X-NH_2_ was added to the cell media for a final concentration of 37 μM per peptide. Each Library was tested in three times, each in triplicate with three technical repeats for a total of 27 wells. 40 μl samples were collected from the media at hours 0, 1, 4, 8, 24, and 48 in duplicate.

In-between timepoints, one set of samples was loaded directly within the LC-MS and measured as soon as possible, then stored at 4°C. The second set of samples was immediately frozen at -80°C. After 48 hours, samples were kept frozen and thawed just prior to LC-MS, and 4μL of acetic acid was added to each well. After three weeks the samples were tested again. A non-degradable internal standard, NH2-βF(βA)6-Amide, where βA is a β-alanine and βF is β-homophenylalanine, was included at a 37 μM concentration for all studies.

### 2.4 Peptide calibration curve studies

Human mesenchymal stem cells (hMSCs) (Rooster Bio, LOT 310368) at passage 3 were seeded into 24 well plates at a seeding density of 75,000 cells per well in 1mL of RoosterBasal-MSC-CC (RoosterBio, SU-022) containing RoosterBooster-MSC (RoosterBio SU-003). After 24 hour the media was changed, and peptide library AcβA-X-RGEFV-NH_2_ was added to the cell media at the following concentrations: 7.4, 14.8, 22.2, 29.6, or 37 μM per peptide. Each concentration was tested in three times, each in triplicate with three technical repeats for a total of 27 wells. 40 μl samples were collected from the media at hours 0, 1, 4, 8, 24, and 48 in duplicate.

### 2.5 Sample injection solvent studies

The AcβA-X-RGEFV-βA peptide library was added to solvents: Water, 50% Acetonitrile in Water, PBS, RoosterBasal-MSC-CC (RoosterBio, SU-022) containing RoosterBooster-MSC supplement (RoosterBio SU-003), VascuLife Basal Media (Llifeline Cell Technology™, LM-0002), and macrophage serum free media with L-Glutamine (Gibco 12065-074) with 1% antibiotic-antimycotic (Gibco, 15240-062) at a concentration of 37 μM per peptide. This was also repeated with each solvent containing 10% acetic acid at a concentration of 37 μM for a total of 12 different solvents tested. The library within any given solvent was tested three times, each in triplicate with three technical repeats for a total of 27 wells. Samples were immediately loaded into the LC-MS for data collection.

### 2.6 Solid phase extraction peptide purification

The samples were then dissolved in 1 mL of water with 0.1% acetic acid and purified using Sep-Pak C18 Plus Short Cartridges. Briefly the columns were conditioned with 6 ml of acetonitrile with 0.1% acetic acid. Once the 0.1% acetic acid in water had been flushed through, the samples were loaded, and it was washed with water containing 4% acetonitrile and 0.1% acetic acid to wash the salts through. The peptides were then eluted with 50:50 acetonitrile:water and 0.1% acetic acid. Both the liquid that was flushed through (the wash) and the eluted peptides were collected separately.

### 2.7 Quantification of column performance

A new column (ProntoSIL C18 AQ, 120 Å, 3 μm, 2.0 × 50 mm HPLC column, PN 0502F184PS030) was first primed by flushing 0.1% acetic acid in ultrapure water, followed by 0.1% acetic acid in acetonitrile, and last with 0.1% acetic acid in ultrapure water all at a volumetric flow rate of 300 μL/min for 20 minutes. Peptide Library AcBA-X-RGEFV-βA was dissolved in ultrapure water containing 10% acetic acid and was used to track performance of the HPLC column. Conditioned media consisting of high glucose DMEM supplemented with 1% vol/vol anti/anti, 7.5% wt/vol BSA, 0.1% vol/vol L-proline [Sigma-Aldrich]), 1% vol/vol insulin-transferrin-sodium selenite (ITS; Sigma-Aldrich), 0.2% vol/vol-dexamethasone (Sigma-Aldrich), 0.1% vol/vol ascorbic acid, 0.1% vol/vol transforming growth factor-β1 (TGF-β1; Peprotech), and 0.1% vol/vol linoleic acid (Sigma-Aldrich) was used.^27^ After every injection of the peptide library into the HPLC, the fouling agent was injected four times. This was continued until column failure was observed via LC-MS.

### 2.8 LC-MS data acquisition

From each sample, 10 μL of crude solution was introduced by the LC-MS through an Thermo Scientific Vanquish LC System (Thermo Fisher Scientific) which outputted to a Thermo Scientific LTQ XL Linear Ion Trap Mass Spectrometer (Thermo Fisher Scientific). The sampled mixture was trapped on a column (ProntoSIL C18 AQ, 120 Å, 3 μm, 2.0 × 50 mm HPLC Column, PN 0502F184PS030, MAC-MOD Analytical Inc.). The samples were loaded onto the column with a solvent containing acetonitrile/water, 5:95 (v/v) containing 1% acetic acid at a flow rate of 300 μL/min and held for one minute. The sample was then eluted from the column with a linear gradient of 5-40% Solvent B (1% acetic acid in acetonitrile) at the same flow rate for five minutes. This was followed by a 1 min ramp up to 100% solvent B, where it was re-equilibrated with solvent A (1% acetic acid) to 5% solvent B over the course of 1 min and held there for 2 min. The column temperature was a constant 29 °C. The mass spectrometer was operated in positive ion mode. Using a heated ESI, the source voltage was set to 4.1 kV, and the capillary temperature was 350 °C.

### 2.9 LC-MS Analysis

Data analysis was performed on Xcalibur Software (Thermo Scientific). Peptides were identified automatically using the Thermo Xcalibur Processing Setup window where the mass (m/z) ±0.5 AMU and expected retention times for each quantified peptide were inputted. This was later used to isolate individual peptide species from the total ion count (TIC) trace using the Thermo Xcalibur Quan Setup window, where the area under the curve was calculated and visually inspected for accuracy. All peptides were normalized to a non-changing internal standard, NH_2_-βF(βA)_6_-Amide, where βA is a β-alanine and βF is β-homophenylalanine. The integrated peak area of the peptide of interest was divided by the NH_2_-βF(βA)_6_-Amide internal standard to create an area ratio. Relative amounts of a peptide of peptide were then calculated by normalized all values to their corresponding time zero area ratio. Calculated data was visualized using RStudio.

### 2.10 Statistical analysis

All statistical analysis was done in R-studio. Data with two conditions analyzed with a Student’s t-test, and data with more than two conditions was analyzed using an ANOVA followed by a Tukey’s post-hoc test.

## 3 RESULTS AND DISCUSSION

### 3.1 Identification of Non-degradable Internal Standards for Peptides

Peptide degradation during culture was assessed by measuring peptide concentrations at different time points and comparing them to the initial levels. LCMS is often employed for quantifying the concentration of biological compounds by injecting a defined sample volume onto a column, followed by chromatographic separation and mass spectrometry. However, a variety of factors can influence the concentration of peptides within the samples independently of peptide degradation. This includes sample evaporation during culture, which increases the concentration of all solutes, dilution during sample processing, or errors in the amount of peptide injected due to factors such as air bubbles. Furthermore, the amount of peptide that is measured on the mass spectrometer can vary due to changes in instrument parameters over time.^7^ To compensate for this variability, quantitative studies in LC-MS typically benchmark each of the analytes to non-degradable internal standards.^28^ Internal standard selection is important to ensure accurate and reliable analysis of LC-MS based quantification of peptide based systems to compensate for variability in external factors on the LC-MS.^29,30^ To that end, a systematic approach was taken to choosing and assessing the appropriateness of the standard based on the requirements that it is stable throughout culture and is well retained on the chromatography column.

Internal standards should have physiochemical properties akin to the analyte(s) of interest, ensuring similar behavior during purification and analysis, while being resistant to degradation during the experiment. A polypeptide is an ideal internal standard for peptide degradation studies, however the susceptibility of peptides to enzymatic degradation was a significant concern. β-amino acids are non-canonical amino acids that have two carbons between amide bonds instead of one and have been shown to be resistant to most proteolytic degradation.^31-33^ With this in mind, we synthesized a series of ten β-alanine-based peptides as candidate internal standards (Table 1). β-alanine (βA) was chosen because it has a chemical structure similar to the amino acid glycine, and the Fmoc-protected β-alanine used during synthesis is inexpensive. Internal standards must not only resist degradation but also remain on the chromatography column and ionize effectively in the mass spectrometer for accurate quantification. The ten internal standards that each contained six β-alanine residues. These molecules quantified the influence of having ionizable N-terminal amines (NH_2_) versus N-terminal acetylation (Ac), the influence of the incorporation of the canonical amino acid phenylalanine (F) or the aromatic β-amino acid β-homophenylalanine (βF) and placing the hydrophobic residues at the end or the center of the internal standard. Each standard was evaluated for retention time on the column and having a consistent amount of peptide present during incubation with cells. These ten peptides were split into two different pooled libraries that were added to cell culture media to enable benchmarking to each other and cultured with human mesenchymal stem cells (hMSCs) for 48 hours.

**Table 1.** A simple linear regression was fit to each of the peptides within two different libraries. Every peptide has an r-squared value close to one, indicating the linear relationship between the measured and actual amounts in MS.

| AcβA-RGEFV-X-βA | | | | | | | | | |
| --- | --- | --- | --- | --- | --- | --- | --- | --- | --- |
| Amino Acid | A | D | E | F | G | H | I/L | K | M |
| R<sup>2</sup> value | 0.981 | 0.975 | 0.979 | 0.981 | 0.977 | 0.975 | 0.979 | 0.983 | 0.977 |
| Amino Acid | N | P | Q | R | S | T | V | W | Y |
| R<sup>2</sup> value | 0.981 | 0.978 | 0.982 | 0.973 | 0.982 | 0.982 | 0.978 | 0.963 | 0.983 |
| AcβA-X-RGEFV-βA | | | | | | | | | |
| Amino Acid | A | D | E | F | G | H | I/L | K | M |
| R<sup>2</sup> value | 0.998 | 0.998 | 0.999 | 0.999 | 1.000 | 1.000 | 1.000 | 0.997 | 1.000 |
| Amino Acid | N | P | Q | R | S | T | V | W | Y |
| R<sup>2</sup> value | 1.000 | 1.000 | 0.998 | 1.000 | 1.000 | 1.000 | 1.000 | 0.999 | 0.999 |

Our results show that NH_2_-(βA)_6_-Amide had poor affinity to the LC-MS column, eluting before the chromatography gradient begins (Figure 2A). Since salts from cell culture media adversely impact ionization and foul the MS, we send the mobile phase from the first 1.5 minutes of LCMS to waste prior to ionization to prevent salt accumulation in the mass spectrometer. Accordingly, peptides and internal standards which elute before the gradient begins are not suitable for quantification since they will not be analyze by the mass spectrometer. N-terminal acetylation, the addition of a phenylalanine, or β-homophenylalanine improves retention time by increasing they hydrophobicity of the polypeptide, which increases the affinity to the column (Figure 2A). After 48 hours of degradation, three internal standards candidates had both good retention times and less than 10% degradation (Figure 2B): 1) Ac-βA_6_-Amide, 2) NH_2_-βFβA_6_-Amide, and 3) Ac-FβA_6_-Amide. The ability of molecules to be identified in mass spectrometry is dependent on their propensity to become ionized, and the Ac-βA_6_-Amide standard was excluded due to weak signal intensity of the LC-MS (not shown), likely due to the lack of charged groups. NH_2_-βFβA_6_-Amide was chosen for its highly repeatable peptide elution time of 2.68 minutes, its repeatability in measurement within 48 hours of incubation with within the sample matrix, and its net charge of +1 at in the LC-MS mobile phase.

**Figure 2.**
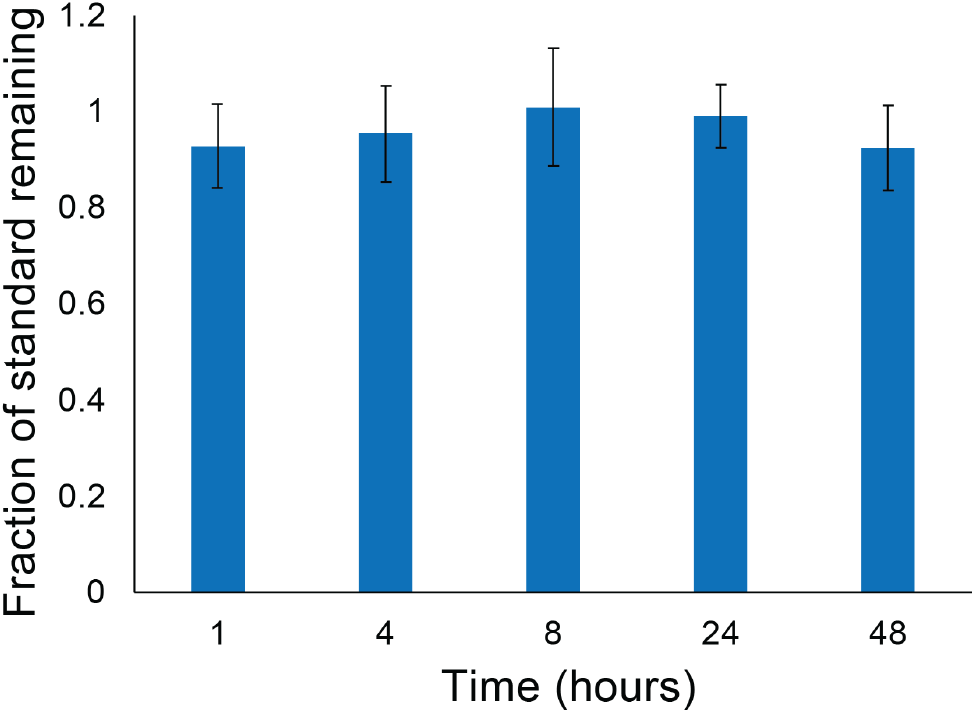
A NH_2_-βFβA_6_-Amide internal standard has physiochemical properties that are similar to peptides, but is stable from proteolytic degradation for 48 hours in cell culture media.

### 3.2 Validation of sample linearity within complex mixtures

Peptide degradation during culture was assessed by measuring peptide concentrations at different time points and comparing them to the initial levels. LC-MS is often employed for quantifying the concentration of biological compounds by injecting a defined sample volume onto a column, followed by chromatographic separation and mass spectrometry. However, a variety of factors can influence the concentration of peptides with in the samples independently of peptide degradation. This includes sample evaporation during culture, which increases the concentration of all solutes, dilution during sample processing, or errors in the amount of peptide injected due to factors such as air bubbles. Furthermore, the amount of peptide that is measured on the mass spectrometer can vary due to changes in instrument parameters over time.^7^ To compensate for this variability, quantitative studies in LC-MS typically benchmark each of the analytes to non-degradable internal standards.^28^ Internal standard selection is important to ensure accurate and reliable analysis of LC-MS based quantification of peptide based systems to compensate for variability in external factors on the LC-MS.^29,30^

Internal standards should have physiochemical properties akin to the analyte(s) of interest, ensuring similar behavior during purification and analysis, while being resistant to degradation during the experiment. A polypeptide is an ideal internal standard for peptide degradation studies, however the susceptibility of peptides to enzymatic degradation was a significant concern. β-amino acids are non-canonical amino acids that have two carbons between amide bonds instead of one and have been shown to be resistant to most proteolytic degradation.^31-33^ In this work we used NH_2_-βFβA_6_-Amide as an internal standard, where βF is βA is β-phenylalanine and βA is β-alanine, because the polyamide structure is similar to peptides, but the presence of two carbons between the amide bonds makes these less susceptible to degradation. We found that the NH_2_-βFβA_6_-Amide internal standard was stable in cell culture for at least 48 hours, with a peak integration area being greater than 92% of the peak at 0 hours.

A central goal of this work is to identify a simple methodology that requires minimal effort in obtaining quantitative degradation results from complex mixtures. Standard curves are widely used to convert measured values to absolute amounts of analyte, however generating the standard curves can be laborious, especially for multiplexed assays containing numerous analytes. Ideally there would be a linear dependence of the measured peptide amount in MS compared to amount of peptide present in the solution over the concentration range used in degradation studies. We synthesized two libraries each containing 19 peptides with arange of charge states and hydrophobicities to ensure that the results are robust and not specific to an individual peptide. These libraries have the form AcβA-RGEFV-X-βA and AcβA-X-RGEFV-βA, where AcβA is an acetylated β-alanine on the N-terminus, and the position X contains peptides with every canonical amino acid except cysteine, which forms disulfide bonds under physiological conditions. The two different libraries have the variable amino acid position on either the N-terminus or C-terminus to account for any influence of variable chemistry at each terminus. We made a series of dilutions of the peptide libraries that kept the concentration of the NH_2_-βFβA_6_-Amide internal standard constant and varied the levels of analyte libraries (Figure 3). Starting with an initial concentration in which each of the peptides in the libraries had a 37 micromolar concentration, we performed dilutions with 80%, 60%, 40%, and 20% of the initial peptide concentration. A graph containing a representative subset of the data indicates that the relationship between the amount in the sample and calculated amount in mass spectrometry was linear for all peptides, typically having an R-squared value above 0.98, indicating a high degree of correlation (Figure 3). Table 1 details the R-squared values for all samples, showing that while the Ac β A-X-RGEFV-β A peptides tended to have higher R-squared values than AcβA-RGEFV-X-βA peptides, all the tested peptides had a high degree linearity.

**Figure 3.**
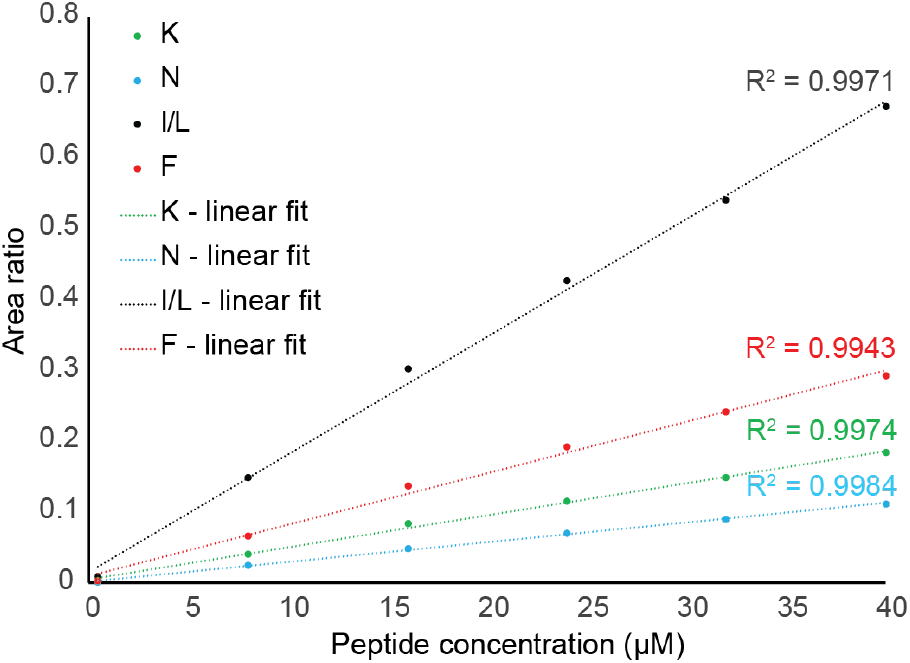
Calibration curves for a subset of the AcβA-X-RGEFV-βA peptide library show that the linear relationship between the measured amount on the LC-MS and the concentration of peptide in the sample. The area ratio is the ratio of the area of the peptide peak in mass spectroscopy divided by the area of the internal standard peak.

### 3.3 Effect of Sample Storage on Peptide Stability

In our degradation studies we incubate peptide libraries with cells for a defined period of time, after which peptide degradation is quantified using LC-MS. Sample injection onto the LC-MS is not typically done immediately after culture, and in most cases the samples are stored in a -80°C freezer prior to analysis. Before analysis the samples are thawed and the plates are loaded into the LC-MS, which is kept at 4°C, however for extended runs an individual sample may not be injected onto the LC-MS for more than 72 hours. The collected cell culture media contains proteases, such as matrix metalloproteinases^34^ and others that have the potential to further degrade peptide libraries after the sample has been collected, which is undesirable. The peptide library AcβA-RGEFV-X-βA was previously shown to degrade in the presence of hMSCs over a 48-hour period and we used this library to quantify the effects that sample storage has on further peptide degradation. The overall goal of these experiments was to identify simple storage conditions that minimize sample degradation over a three-week storage period.

To quantify the effects of storage conditions on sample quality, 40 μL of cell culture media was taken and a portion was immediately injected onto the LC-MS to quantify the amount of peptide initially present in the samples. The plates in the untreated condition were then stored at 4°C and treated samples were frozen at -80°C until after the 48-hour timepoint, where they were treated with 4μL of acetic acid, injected onto the LC-MS, and then stored again at -80°C. It is worth noting that the untreated and acidified data is pared, with both being draw from the sample at the same time. After 23 days the samples were injected again onto the LC-MS and it was seen that the acetic acid treated samples had minimal variability within replicates, as quantified using the coefficient of variation, which divides the standard deviation of the replicates by the average value for the replicates. The untreated samples, however, showed more statistically significant variability after 24 hours across all amino acids in the library (Figure 4).

**Figure 4.**
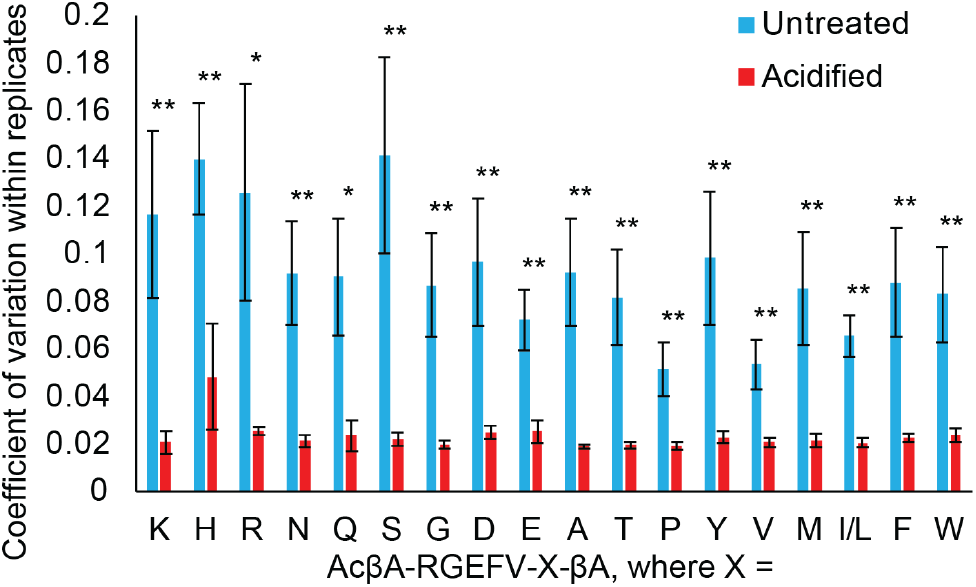
Samples were either treated with acetic acid, or left untreated and stored at 4*°*C, while the acidified samples were stored at -80*°*C and acidified to 10% acetic acid prior to LC-MS. We then ran the samples on LC-MS and found that storing the samples at -80°C and treating them with acetic acid reduced variability within technical replicates.

### 3.4 Effects of Sample Solvent Conditions on Chromatography

Reverse-phased chromatography is a highly effective technique for separating peptides.^35^ The use of a hydrophobic sorbent as the stationary phase results in affinity-based partitions, determined by the interactions of the solute with the mobile phase and the stationary phase.^35^ Ideally samples should be dissolved in solvents having the same composition as the mobile phase that is present within the system when the sample is being loaded onto the column. However, this can require processing steps to remove the initial solvents in which the samples are dissolved, and replacing them with the mobile phase, which can be both time consuming and increases the chances of sample loss.

To study effects of initial peptide solvent on column performance, the peptide library AcβA-X-RGEFV-βA was chosen, where X is every amino acid except cysteine. Previous work has shown that chemistry of peptide termini is the most effective means by which non-specific degradation of peptides may be controlled as compared to concentration or even PEGylation.^16^ In previous studies, peptides modified with an N-terminal acetylated β-alanine was shown to be the most proteolytically resistant to degradation within a 48-hour period when incubated with a variety of cell types.^16^ This resistance to proteolytic degradation makes the AcβA-X-RGEFV-βA library a good model system to study the effects of the sample matrix on peptide affinity to the stationary phase of the LC-MS, as well as performance on the mass spectrometer.

To better understand the effects that different sample solvent conditions have on chromatography, we dissolved the AcβA-X-RGEFV-βA library in 12 different solvents (Figure 5 and S1). This includes pure water, phosphate buffered saline (PBS), and three different cell culture medias. We also tested conditions with 10% acetic acid to account for the acidification that is used to prevent protease degradation, and acetonitrile, which can deactivate proteases and is used during sample purification. For each of the 12 conditions the peptide libraries were tested in triplicate, each with three technical repeats, for a total of 9 wells per solvent. In previous work salt has been shown to alter relative separation and retention times of species within the sample matrix due to changes in adsorption due to the electrical double layer repulsion.^36^ In our studies the presence of salt in the sample has a minimal effect on the area ratio, but reduces retention time on the stationary phase, particularly for peptides where the X position contains positively charged residues H, K, and R.

**Figure 5.**
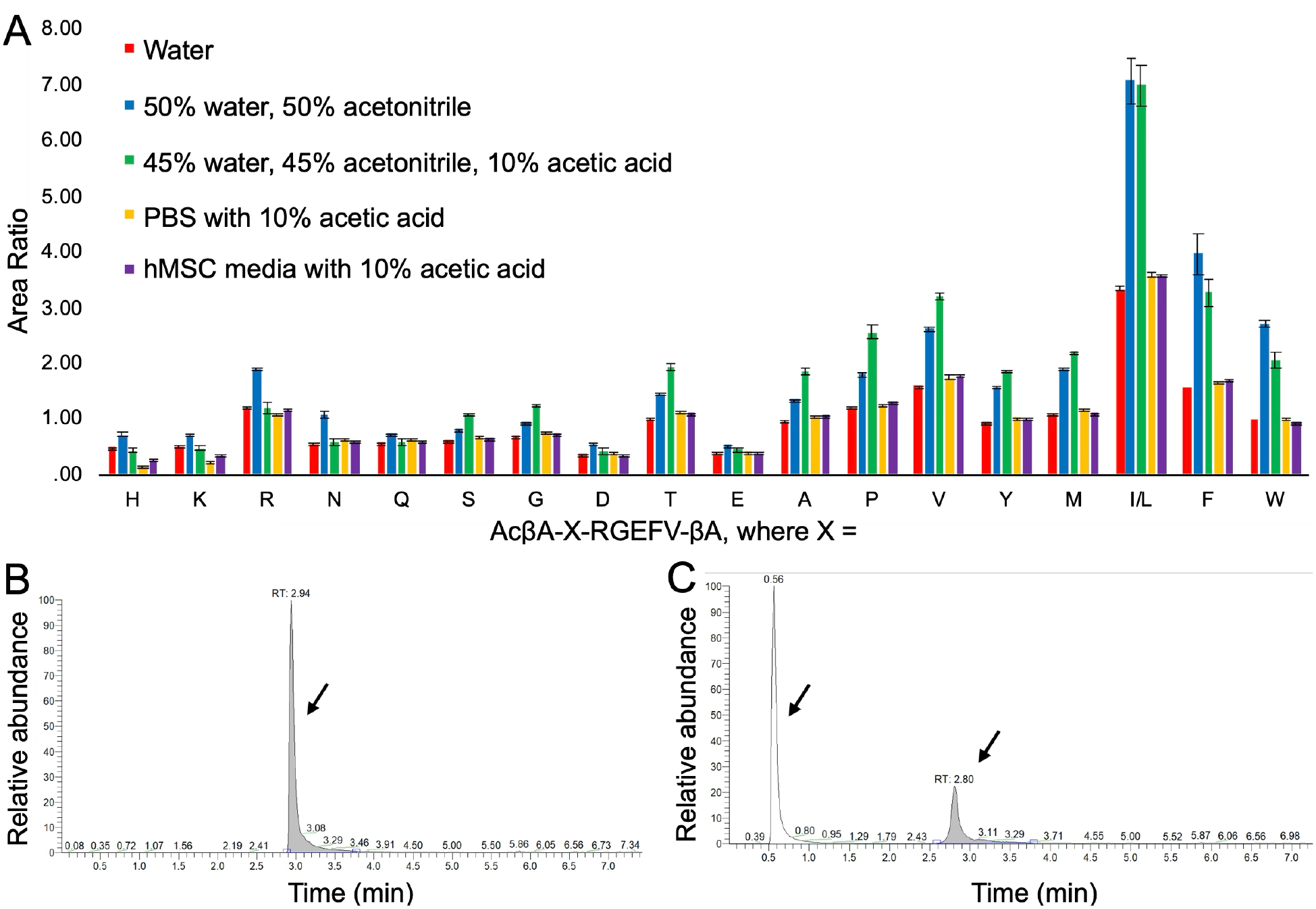
(A) The was dissolved in a variety of common media conditions and solvents, and run on the LC-MS. The area ratio is the ratio of the area of the peptide peak in mass spectroscopy divided by the area of the internal standard peak. The amount of each peptide was quantified, and it was seen that having significant amounts of organic solvents such as acetonitrile, can have a significant effect on the results. This is due to the poor retention on the column, which is most commonly seen in the form of peak splitting. When a peptide is well-adsorbed to the column there is a single peak (B), while those with poor chromatography end up in both the breakthrough peak, which is typically sent to the waste, in addition to the normal peak (C).

We found that presence of acetonitrile in the injected solution had a profound effect on LC-MS performance. Acetonitrile is often mixed with water to improve the solubility of hydrophobic peptides, but we found that 50% acetonitrile in water caused peak splitting in the internal standard such that a significant fraction of the internal standard failed to adsorb to the column and eluted prior to the start of the solvent gradient. Since the mobile phase prior to the start of the gradient is typically sent to the waste, the presence of the early elution of the internal standard will adversely impact the quantification of every analyte within the sample. This is problematic because the effects of acetonitrile on the AcβA-X-RGEFV-βA peptide library was not consistent, and a portion of the hydrophilic peptides also failed to adsorb to the column, while the hydrophobic peptides had good retention across all 12 solvent conditions. It is also notable that the addition of 10% acetic acid to the sample appears to increase column performance after the more hydrophilic residues (Figure 5A).

### 3.5 Quantification of Chromatography Column Failure

A key benefit of liquid chromatography-mass spectrometry is that individual runs are often less than 12 minutes, and sometimes less than 5 minutes. Sample injection and collection is typically automated and hundreds of samples can be loaded at a time, which enables LC-MS instruments to run dozens to hundreds of samples per day. While the combination of high throughput and the powerful characterization capabilities of LC-MS is advantageous, this workflow requires preparation of numerous samples for analysis. The presence of lipids and proteins in biological samples is detrimental to the longevity of reverse phase chromatography columns, and a variety of approaches exist to separate these compounds from the analytes of interest prior to LC-MS. For instance, trichloroacetic acid is frequently mixed with samples to precipitate soluble proteins,^37^ and disposable solid phase extraction (SPE) columns, whose properties are similar LC-MS columns, are used to load the crude sample mixture followed by elution of the desired analytes.^38^

A downside of using SPE columns is that each column is typically single use, and generally cost $3-5 per sample, which can be a significant cost burden when analyzing hundreds of samples per week. More generally, both precipitation-based and SPE-based methods have drawbacks in that they need to be optimized for a specific analyte to minimize sample loss and ensure reproducibility, and this is often not possible for studies utilizing libraries of peptides with significantly different physiochemical properties. We validated this by loading our AcβA-X-RGEFV-βA library onto an SPE column and purified the peptides from the proteins, salts, and lipids using the manufacturers instructions. In this process a sample is loaded onto a disposable column, constituents which have poor interactions with the column, such as salts, are washed through, and then the sample is eluted, leaving chemical species such as proteins and lipids on the column. We then ran LC-MS on both the crude mixture prior to purification and the initial wash waste, and found that all peptides in the library were present in the waste, but that hydrophilic peptides are enriched compared to hydrophobic peptides (Figure 6). This highlights the difficulty in purifying libraries of analytes with different physiochemical properties prior to analysis. Furthermore, performing these purification steps on hundreds of samples is a significant time and experimental burden.

**Figure 6.**
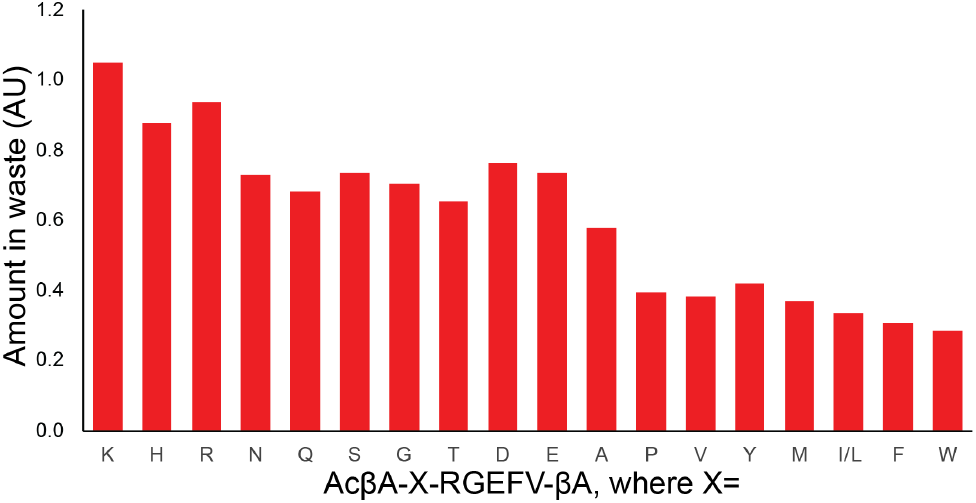
The AcβA-X-RGEFV-βA library was dissolved in water, loaded onto a solid phase extraction column, and then both the peptide in the flow through waste was quantified. Every peptide in the library was found in the waste, however hydrophilic peptides were enriched compared to the hydrophobic sequences. conditions and run on the LC-MS.

We sought to greatly simplify sample preparation by injecting the biological samples directly on the LC column without any purification steps. This prevents any sample losses prior to analysis, requires minimal effect for even for hundreds of samples, and negates the need for costly consumables. A significant downside to this approach is that the proteins and lipids present in biological samples are detrimental to the performance of the LC column. While these columns are not typically treated as disposable, they can be purchased for under $300, which is not prohibitively expensive if each column can be used for hundreds of runs. We quantified changes in the column properties after repeated sample injection by recording changes the column pressure, retention time of each of the analytes on the column, and their integrated area in mass spectrometry to better understand how repeated sample injections influences column performance (Figure 6). It should be noted that specific types of media are used when culturing a cell type. Media formulations frequently incorporate fetal bovine serum (FBS), which is rich in proteins having total concentrations between 30-45 mL/mL.^39^ In these studies, we used media with 10% FBS to quantify the influence that repeated sample injections have on column performance. Due to the variability inherent in FBS, there has been an emphasis on the use of fully defined media formulations.^40^ Fully defined medias typically have lower protein content than those with FBS and peptides dissolved in these medias should reduce column fouling, which will likely increase the number of LCMS samples that can be run before the column needs to be replaced.

To study the effects of repeatedly injecting cell culture samples containing proteins on column lifetime and consistency in data collection, we performed hundreds of injections of media containing 7.5% FBS. During these runs we ran the AcβA-X-RGEFV-βA peptide library every fifth injection to quantify column performance over time, including sample and column characteristics. The accumulation of lipids and proteins on the column will physically clog the column and increase the pressure at any given flow rate.^41^ We observed a steady increase in pressure over the first 200 injections, with the pressure increasing by ∼100 bar (Figure 7A). Injections were repeated for over 600 runs, during which time the retention time of each peptide was also found to be consistent (Figure 7B). An exception to this are a few notable instances in which the column was used after days of disuse, where the retention time was evenly shifted across all analytes (black arrows in Figure 7B).

**Figure 7.**
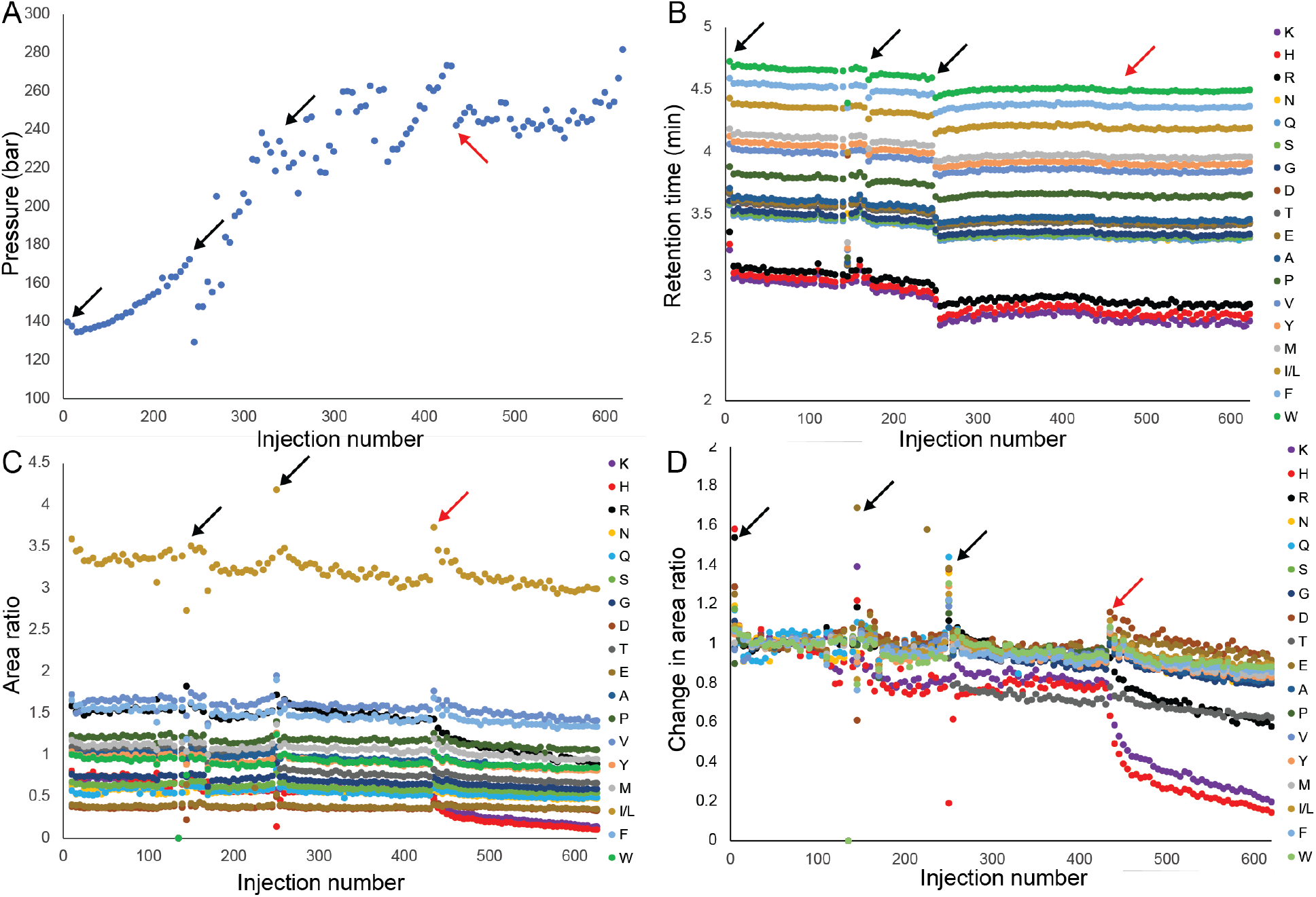
To evaluate the lifespan and performance of LC columns during degradation studies, we conducted a repeated injection protocol totaling hundreds of injections. The protocol involved injecting the AcβA-X-RGEFV-βA peptide library once, followed by four consecutive injections of media with 10% FBS; this entire sequence was repeated 124 times until column failure. The arrows indicate when the first injection was performed after at least 24 hours of non-use. The red arrow marks the onset of column failure where failure of the column is initiated. A) The LC-MS pressure at the beginning of sample runs increases steadily for the first 300 injections and then plateaus. B) The retention time for each analyte decreases slightly over the course of hundreds of injections. C) The area ratio of most peptides, which the area of the analyte divided by the area of the internal standard, decreases slightly over the series of injections, except for hydrophilic molecules. D) Normalizing the area ratio to the values from the initial injections shows that column failure occurs when hydrophilic peptides are no longer retained on the LC column.

In our studies the hallmark of column failure is the inability for hydrophilic peptides to be retained on the column. We started to observe column failure after 348 injections of the media containing 7.5% FBS and 87 injections of the library, for a totally of 435 injections (Figure 7C). To better quantify changes in column performance, we normalized the area ratio of each peptide during the initial runs on the column as a baseline, and then divided all runs by this normalized value (Figure 7D). It should be noted that the first two or three injections of a series of runs, as depicted by the arrows on Figure 7, can have substantially altered chromatographic characteristics compared to subsequent injections, and this effect is independent of column health. We found that after 435 injections the most hydrophilic sequences, which are the AcβA-X-RGEFV-βA peptides containing the positively charged amino acids lysine and arginine in the X position, started showing decreased area ratio due to poor retention on the column. These results show that column failure is typified by the poor retention of hydrophilic peptides, but highlight that using libraries with hydrophilic sequences are effect for assessing column health.

## 5 Conclusions

In this work we have developed a method for quantifying the degradation of peptide libraries within complex biological samples. This method utilizes widely-available instrumentation, and was designed to minimize the need for consumables, while reducing the time consuming preparation steps. Peptides can be easily made on automated synthesizer and show great promise as therapeutics and as components within biomaterials. Library-based peptide assays which quantify peptide degradation under physiological conditions are needed to improve our control over cell-peptide interactions, and this work will help scientists and engineers use functional approaches to improve biomedical therapies and platforms.

## Acknowledgements

We would like to acknowledge our funding sources, the NIH (1R21GM143593-01) and NSF (Award 2138723). We would like to thank the lab of Lesley Chow for use of their preparative high performance liquid chromatography.

