## Supplemental Information for "A Streamlined High-Throughput LC-MS Assay for Quantifying Peptide Degradation in Cell Culture"

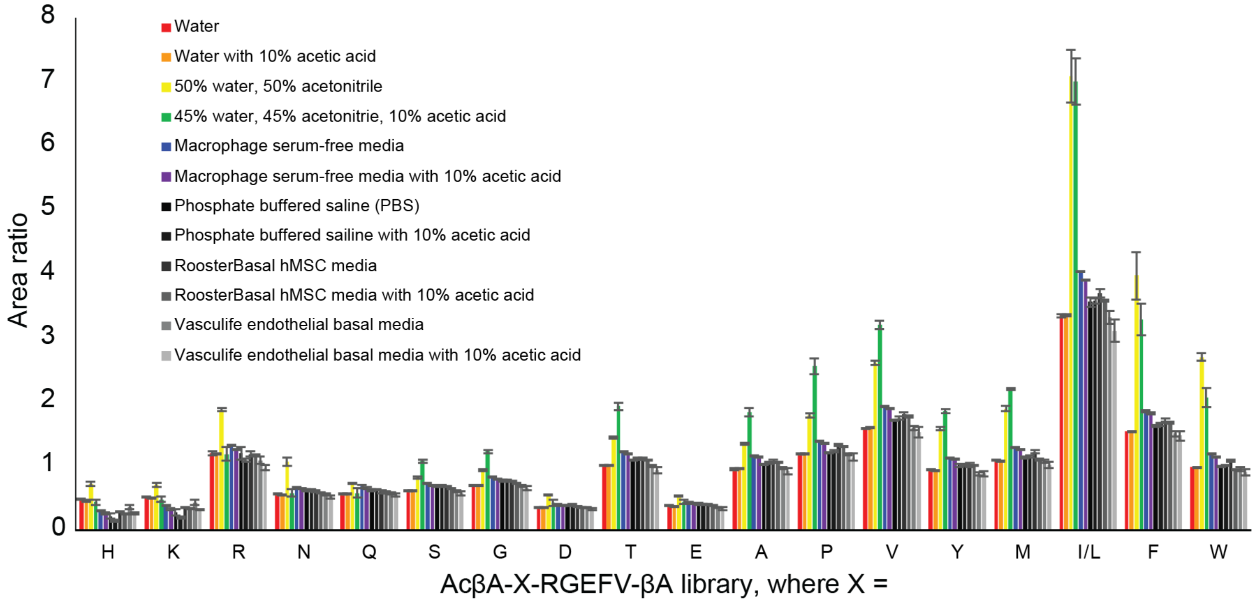


**Figure S1**. Degradation of the AcβA-X-RGEFV-βA peptide library in a variety of solvent conditions.

**A** **Ac-βA-Ala-RGEFV-NH$_2$**

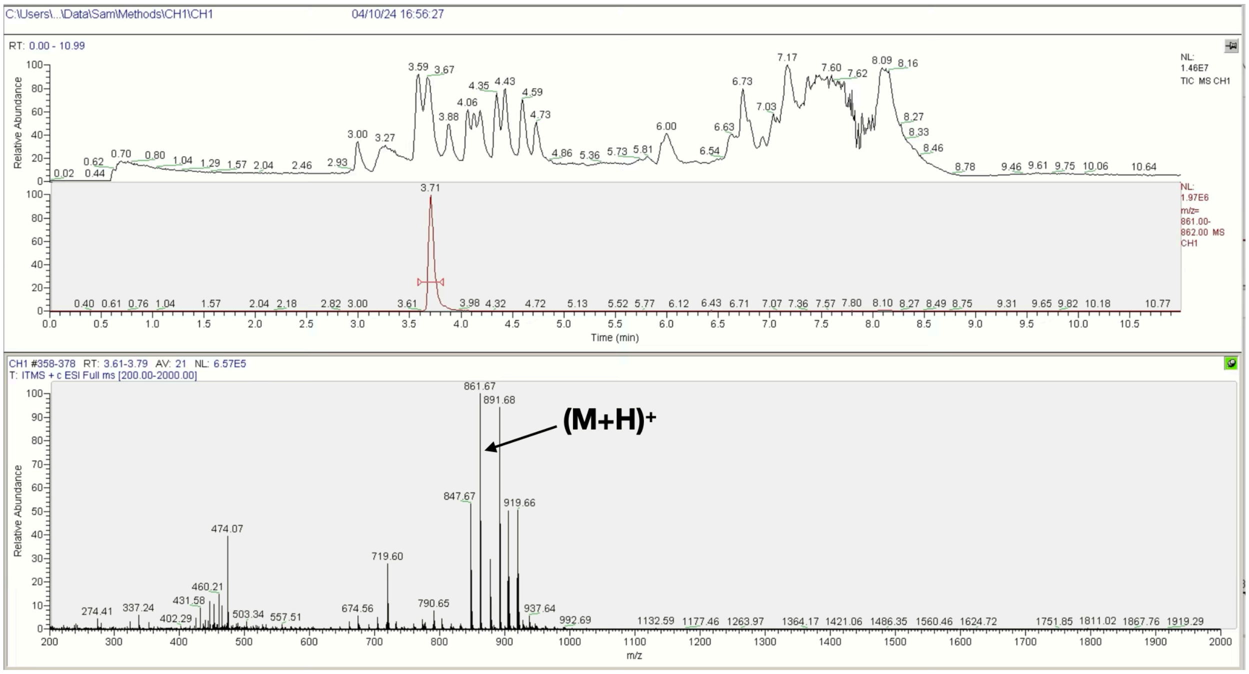


**B** **Ac-βA-Arg-RGEFV-βA-NH$_2$**

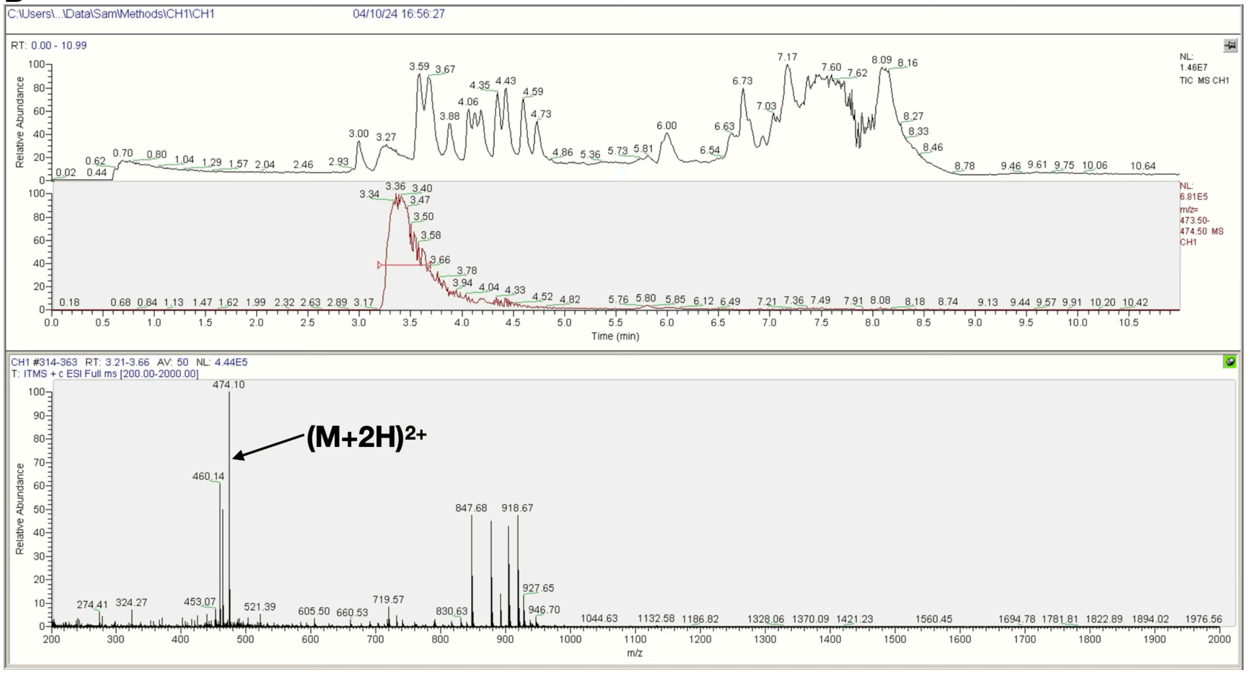

**C**

**Ac-βA-Asn-RGEFV-βA-NH2**

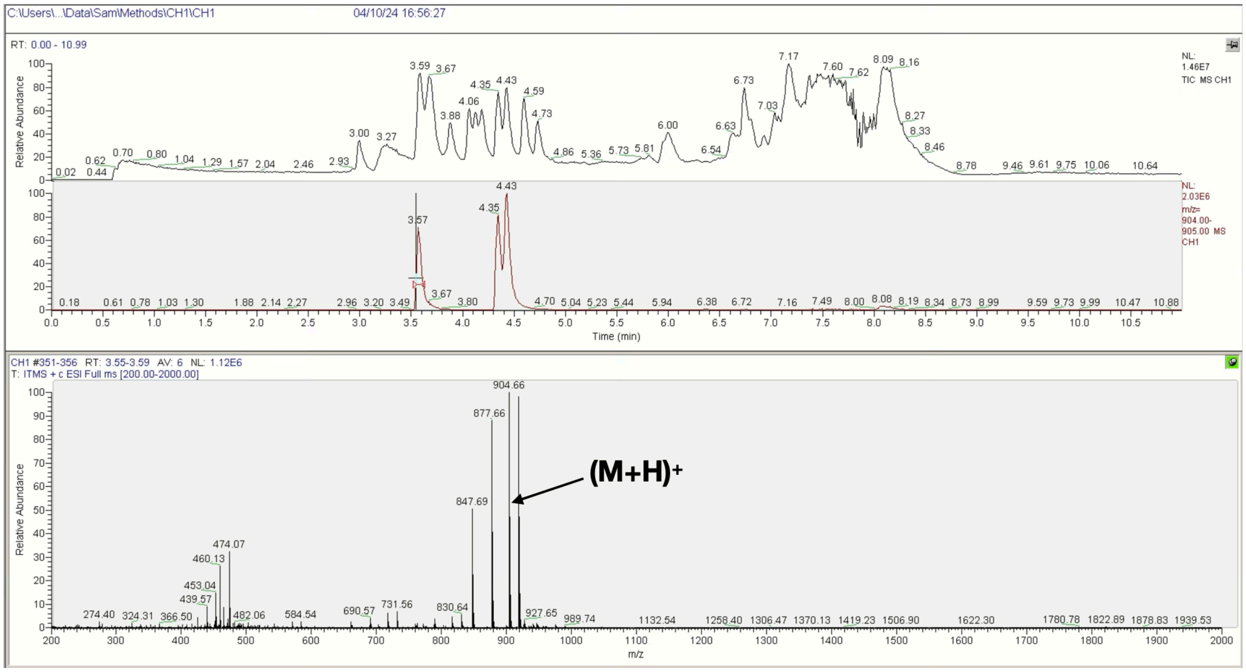


**D**

**Ac-βA-Asp-RGEFV-βA-NH2**

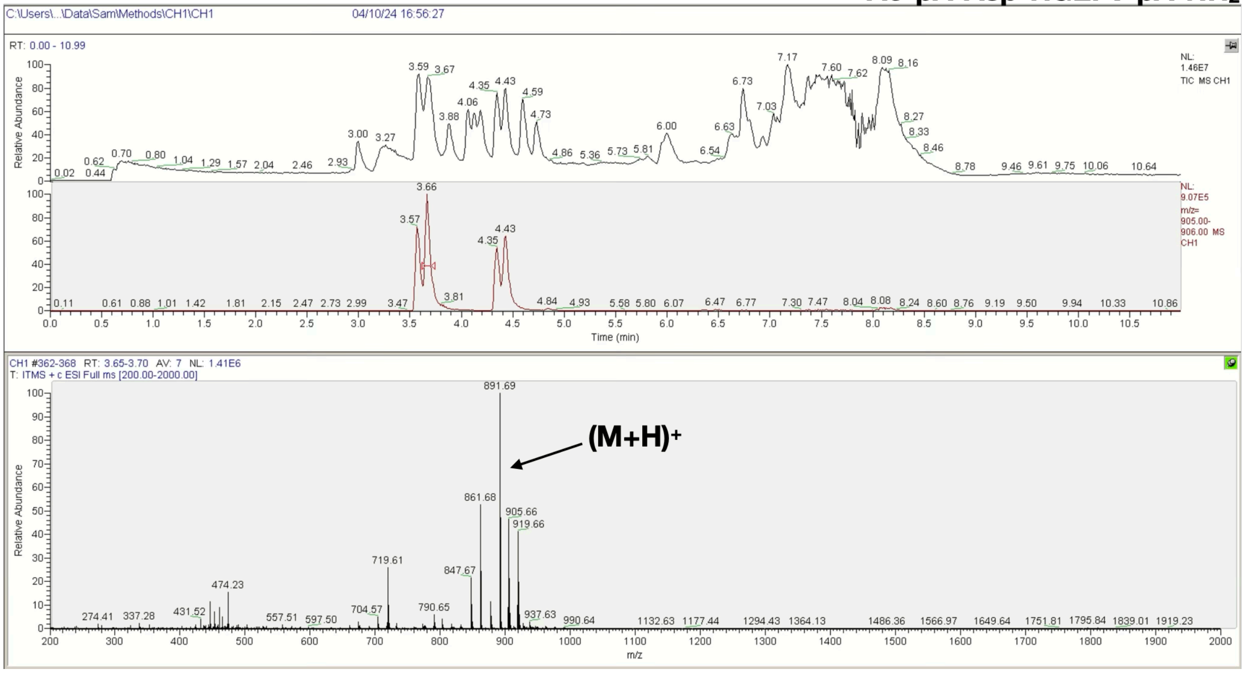

**E** **Ac-βA-Gln-RGEFV-βA-NH$_2$**

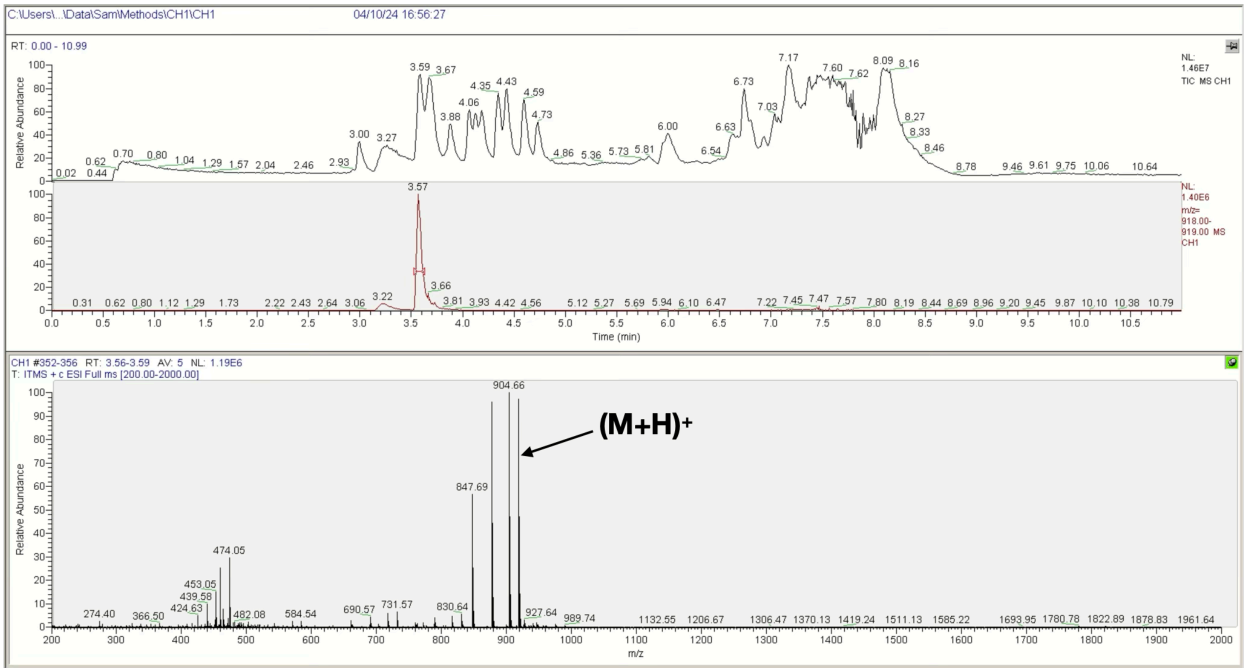


**F** **Ac-βA-Glu-RGEFV-βA-NH$_2$**

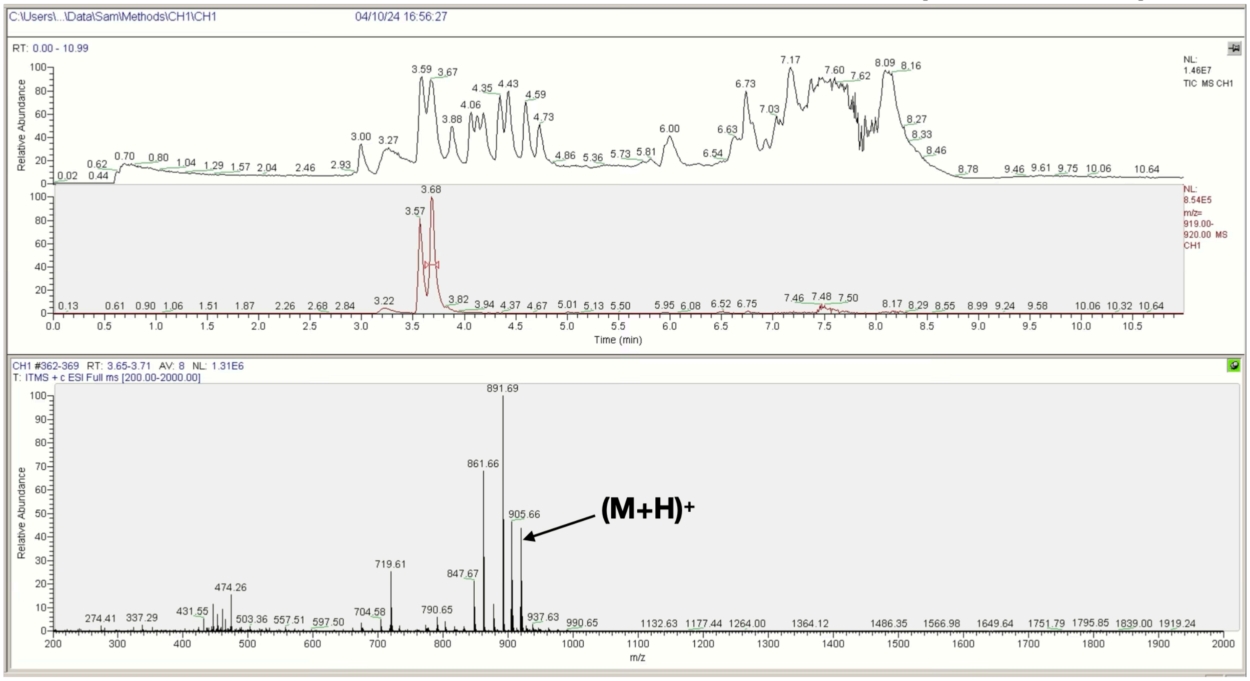

G

**Ac-βA-Gly-RGEFV-βA-NH2**

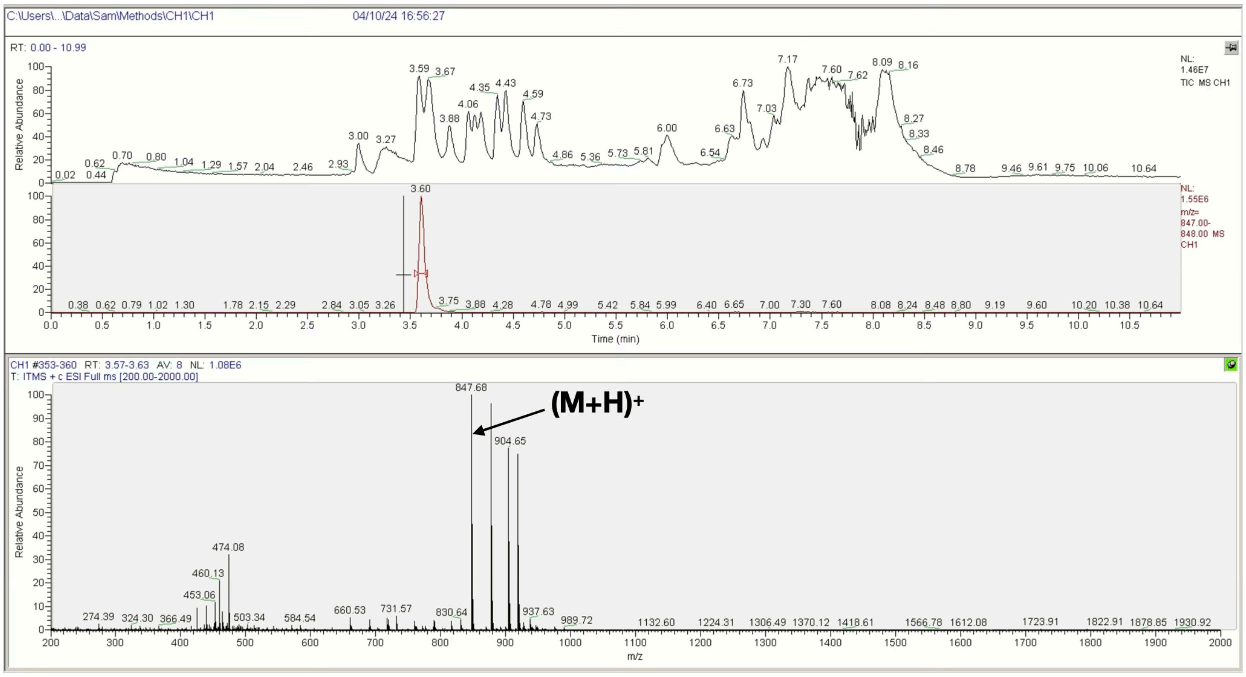


H

**Ac-βA-His-RGEFV-βA-NH2**

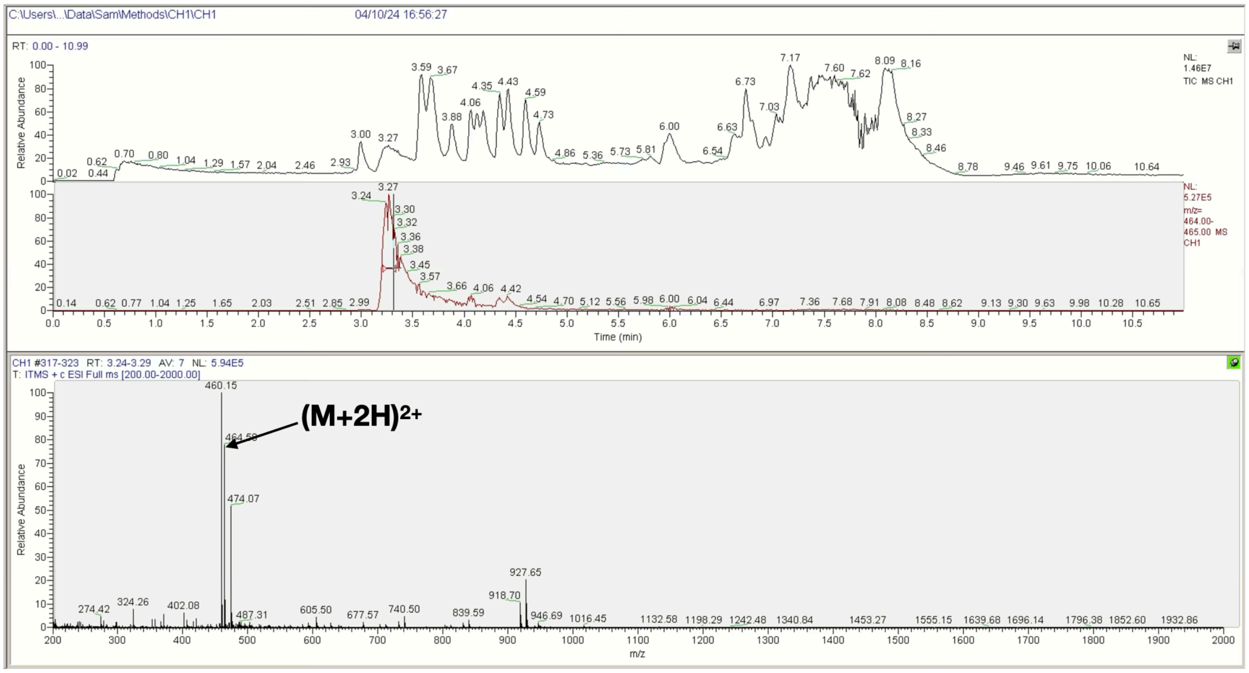

**I** **Ac-βA-Ile/Leu-RGEFV-βA-NH$_2$**

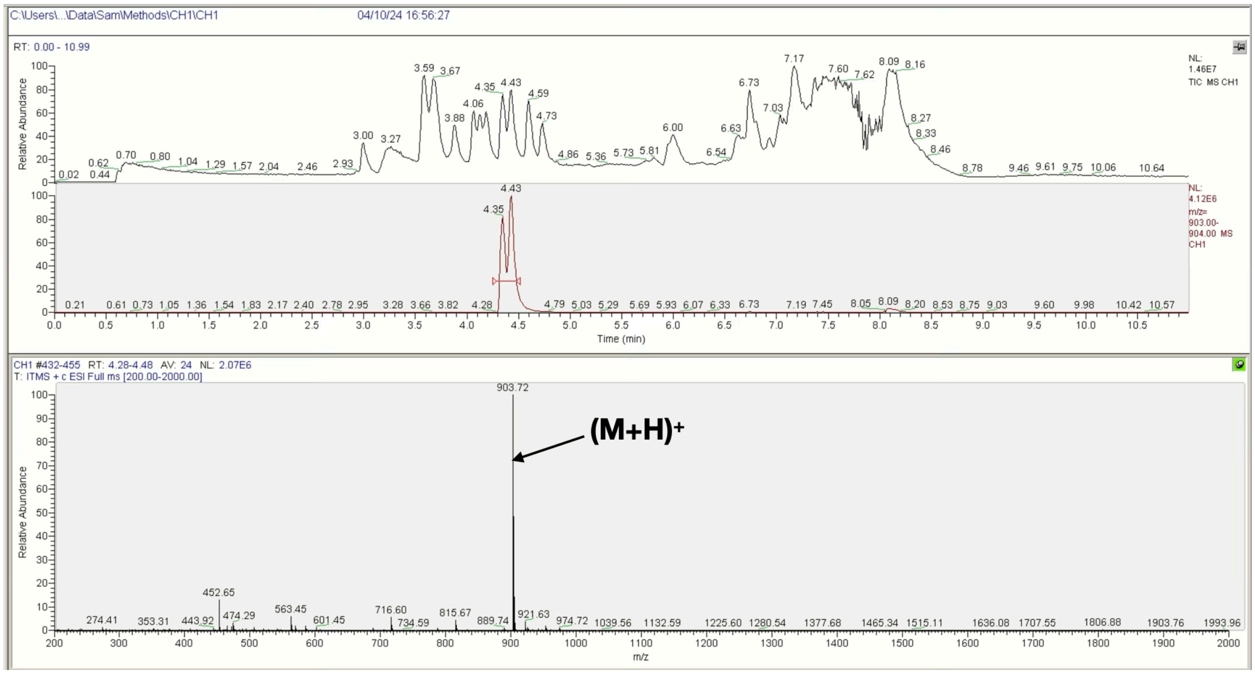


**J** **Ac-βA-Lys-RGEFV-βA-NH$_2$**

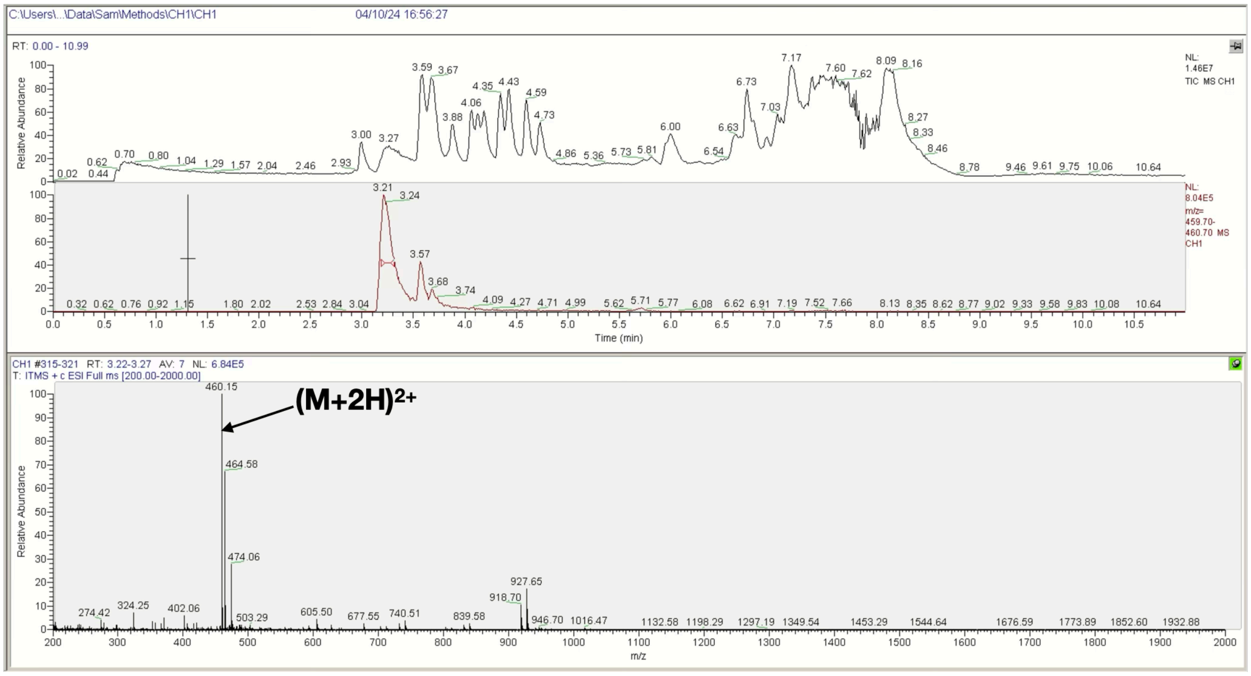

**K** **Ac-βA-Met-RGEFV-βA-NH₂**

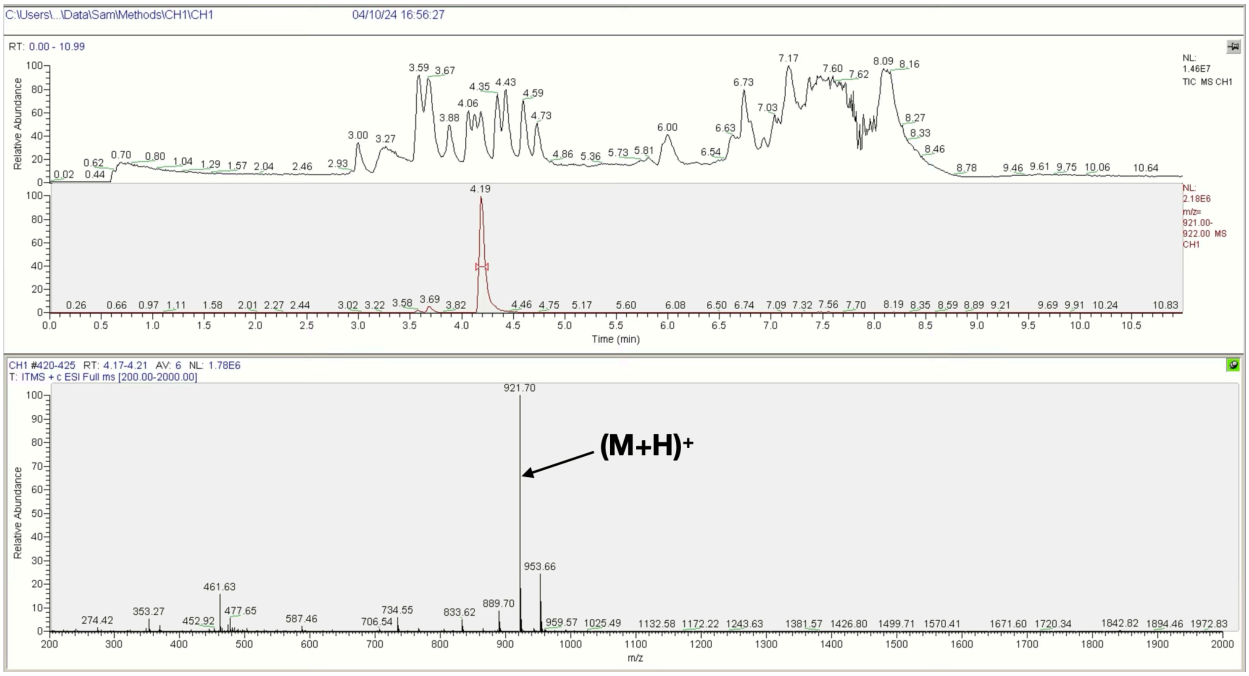


**L** **Ac-βA-Phe-RGEFV-βA-NH₂**

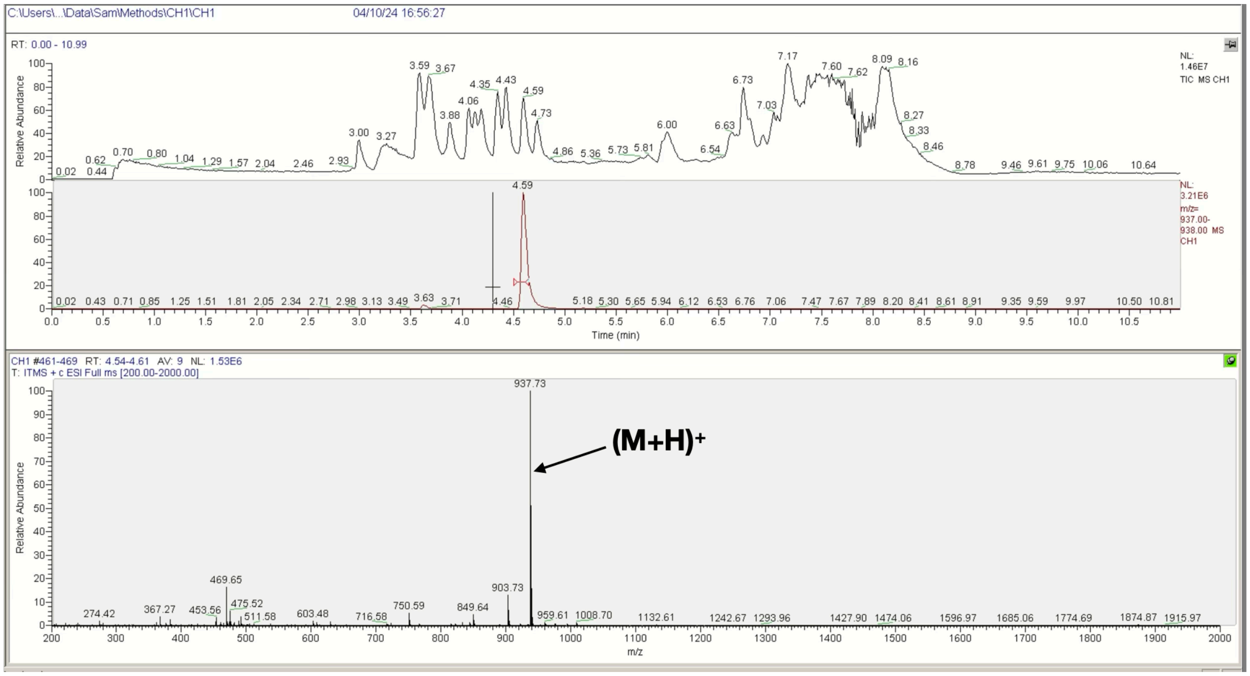

**M** **Ac-βA-Pro-RGEFV-βA-NH₂**

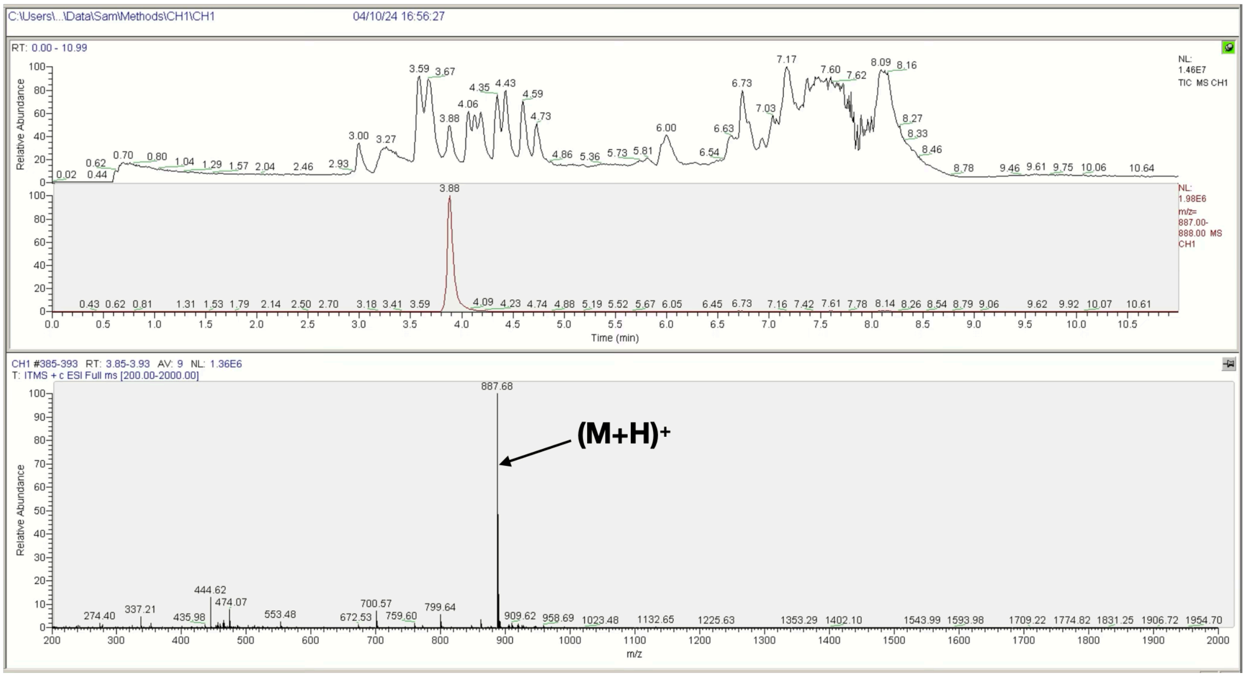


**N** **Ac-βA-Ser-RGEFV-βA-NH₂**

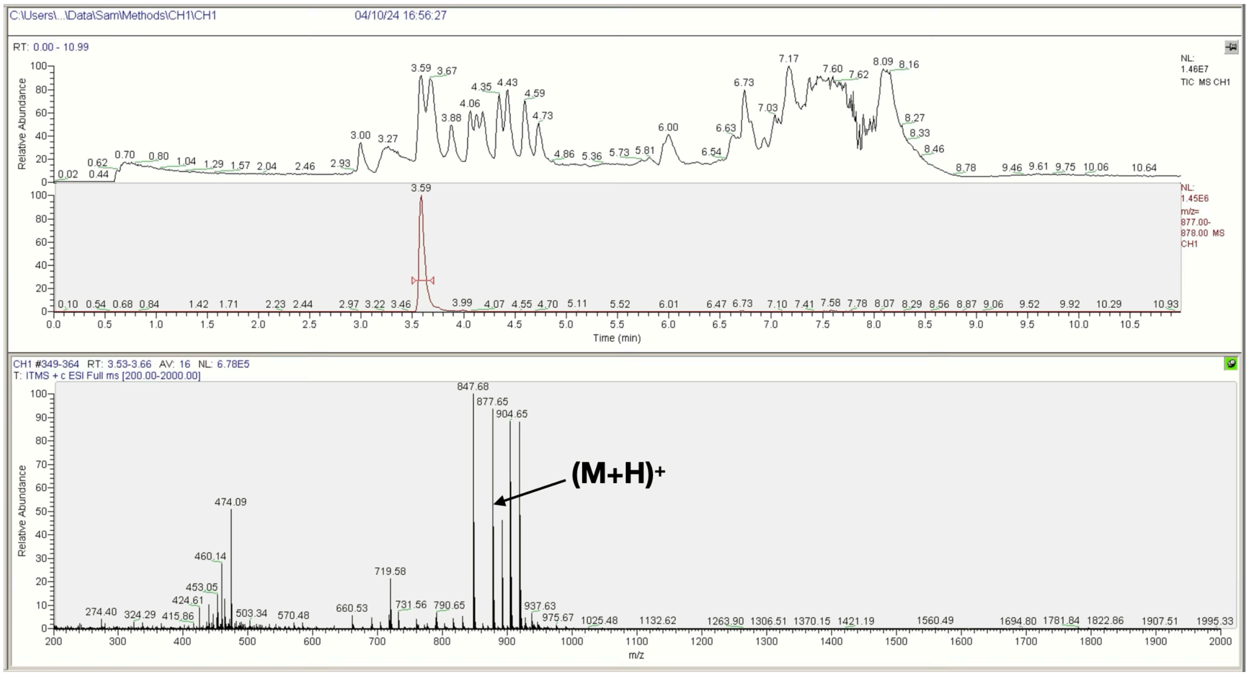

**O** **Ac-βA-Thr-RGEFV-βA-$NH_2$**

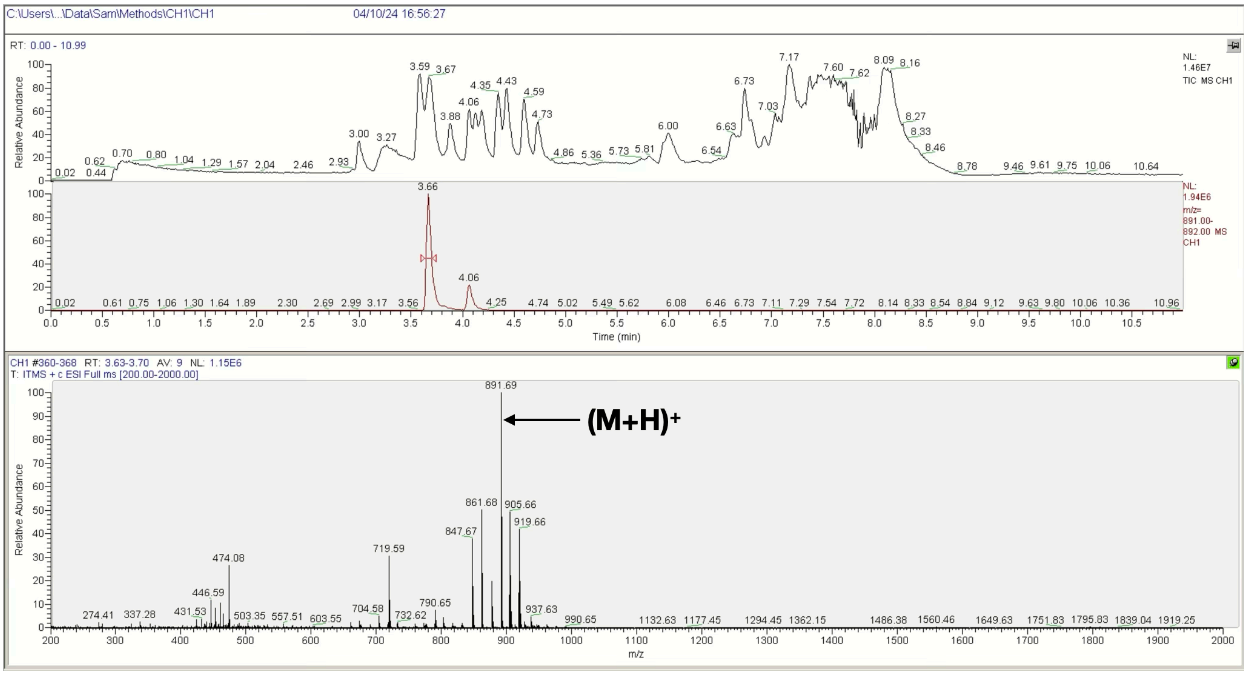


**P** **Ac-βA-Trp-RGEFV-βA-$NH_2$**

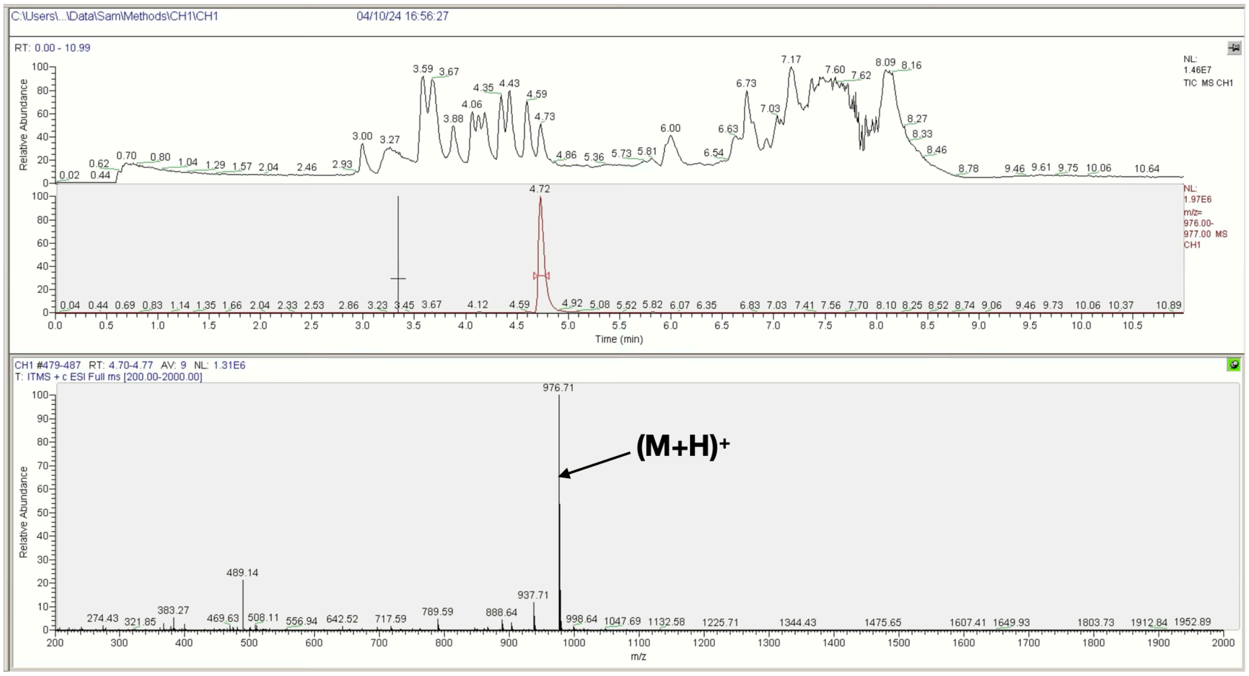

**Q** **Ac-βA-Tyr-RGEFV-βA-NH$_2$**

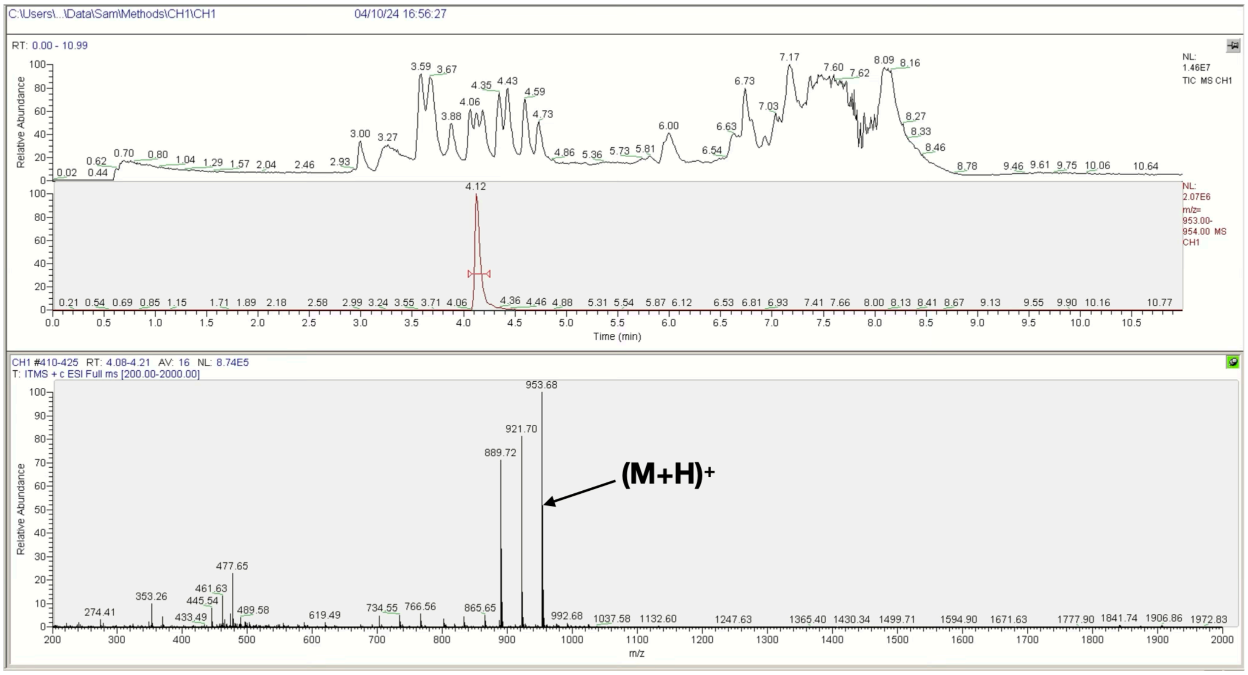


**R** **Ac-βA-Val-RGEFV-βA-NH$_2$**

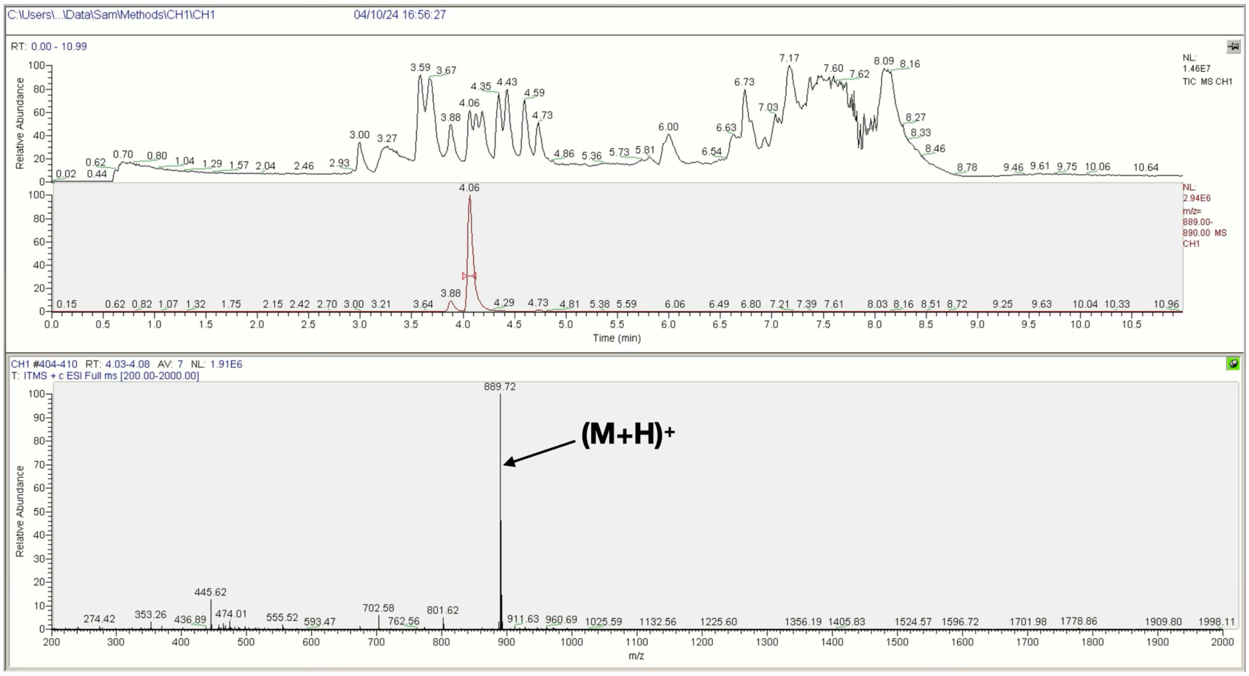


**Figure S2**. LCMS spectra of the Ac-βA-X-RGEFV-NH$_2$ libraries, where X = a) Ala, b) Arg, c) Asn, d) Asp, e) Gln, f) Glu, g) Gly, h) His, i) Ile/Leu, j) Lys, k) Met, l) Phe, m) Pro, n) Ser, o) Thr, p) Trp, q) Tyr, r) Val.

**A** **Ac-βA-RGEFV-Ala-βA-NH₂**

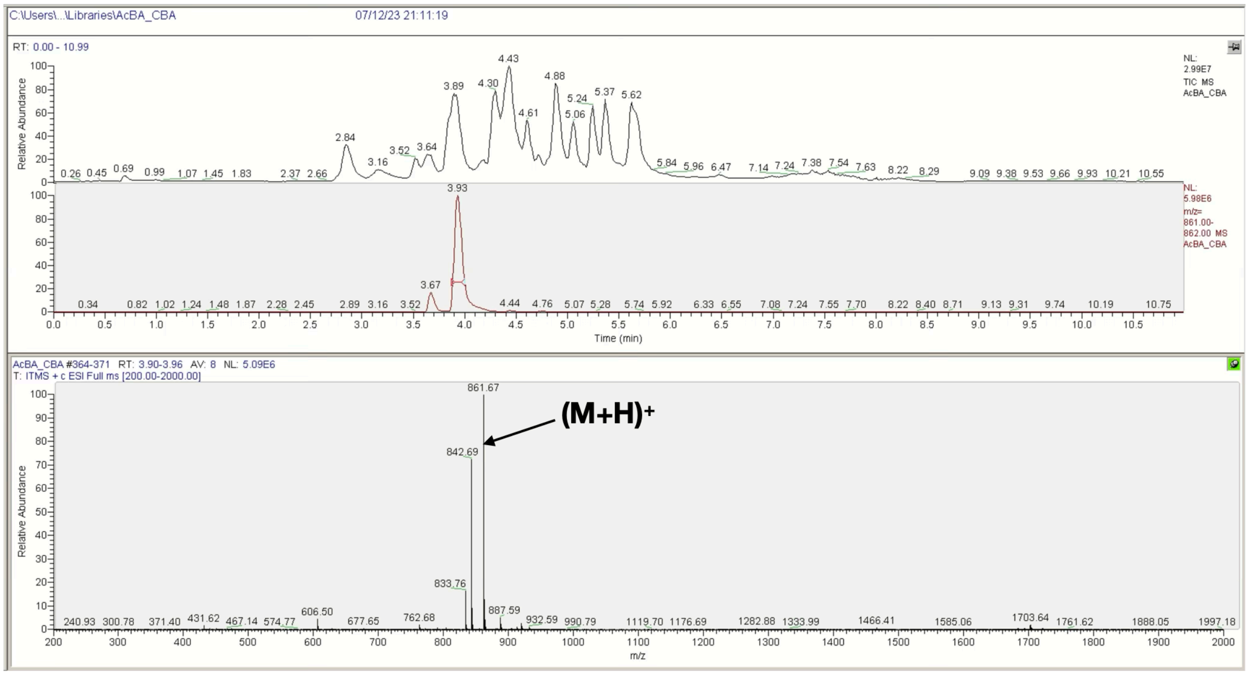


**B** **Ac-βA-RGEFV-Arg-βA-NH₂**

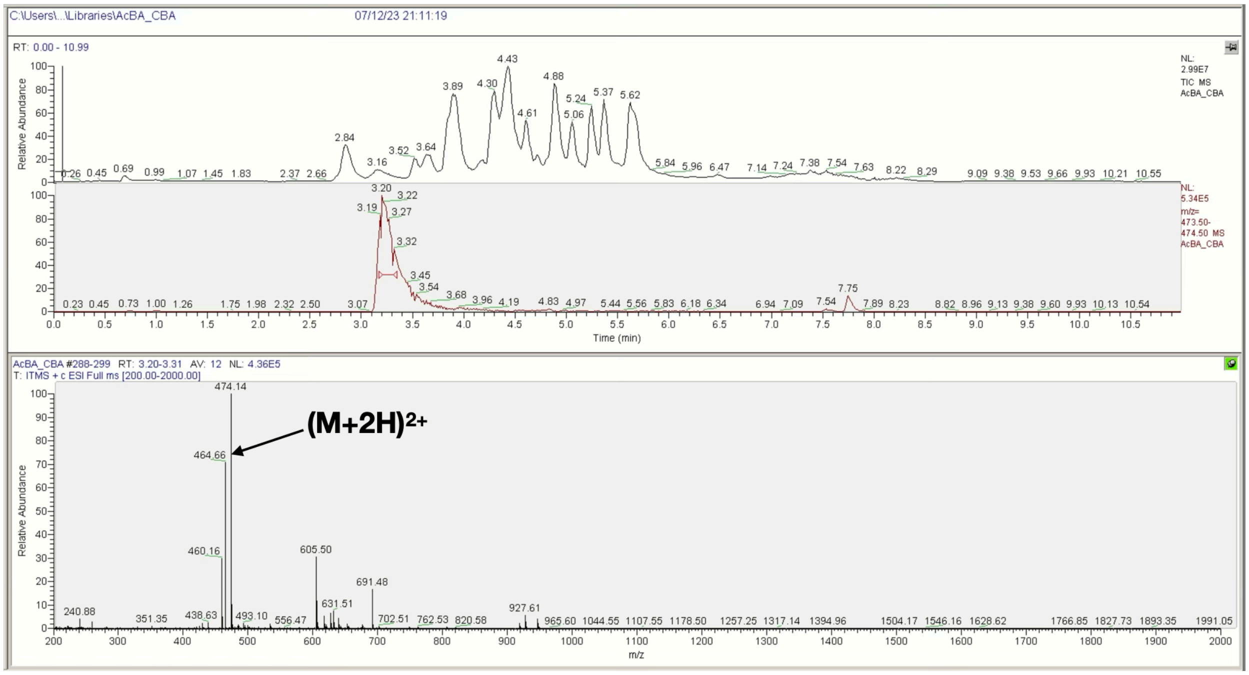

**C** **Ac-βA-RGEFV-Asn-βA-NH₂**

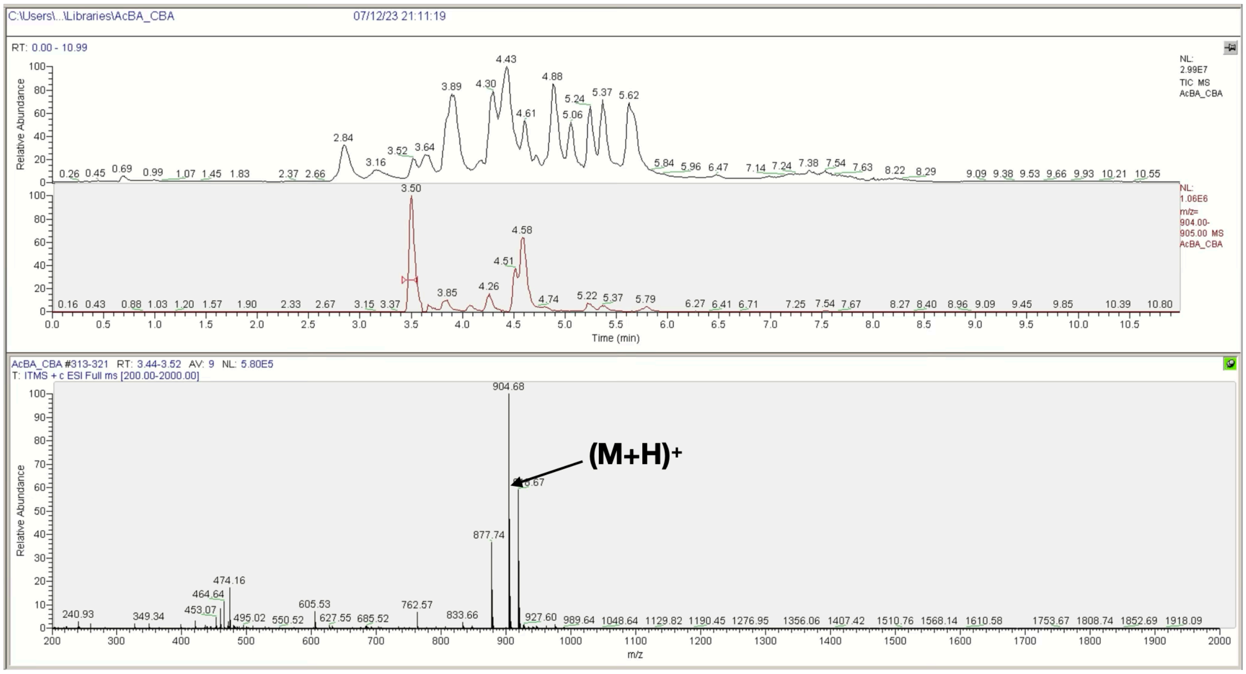


**D** **Ac-βA-RGEFV-Asp-βA-NH₂**

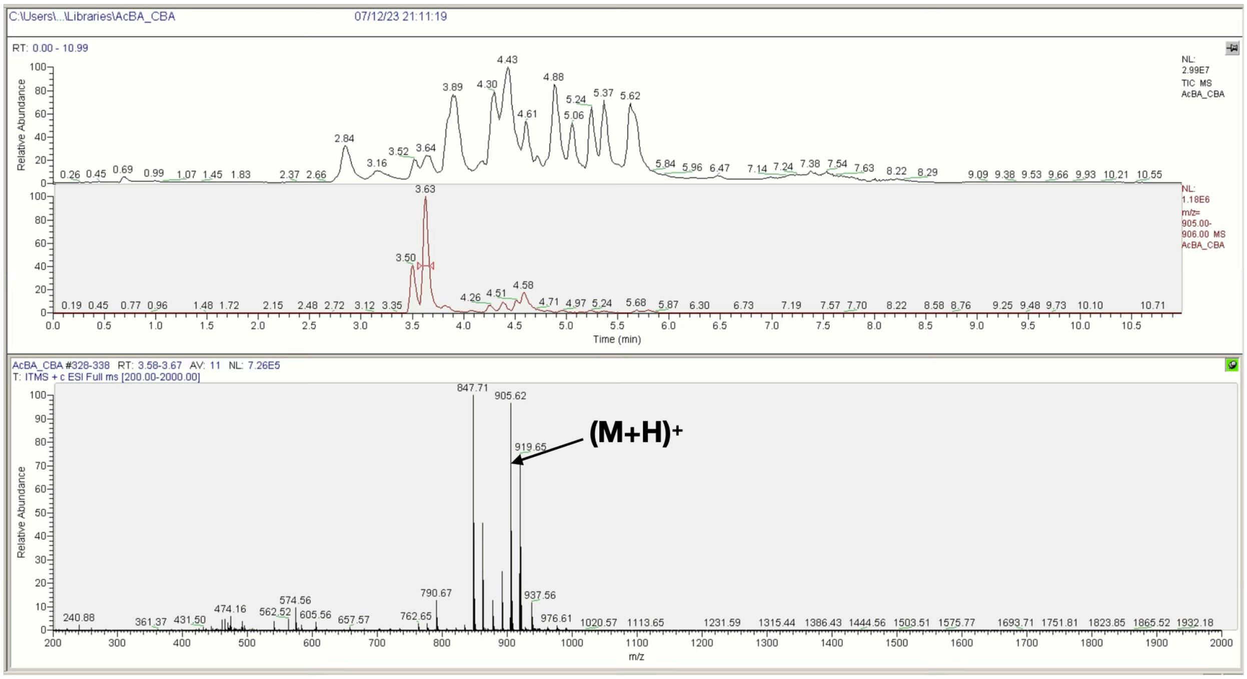

**E** **Ac-βA-RGEFV-Gln-βA-NH2**

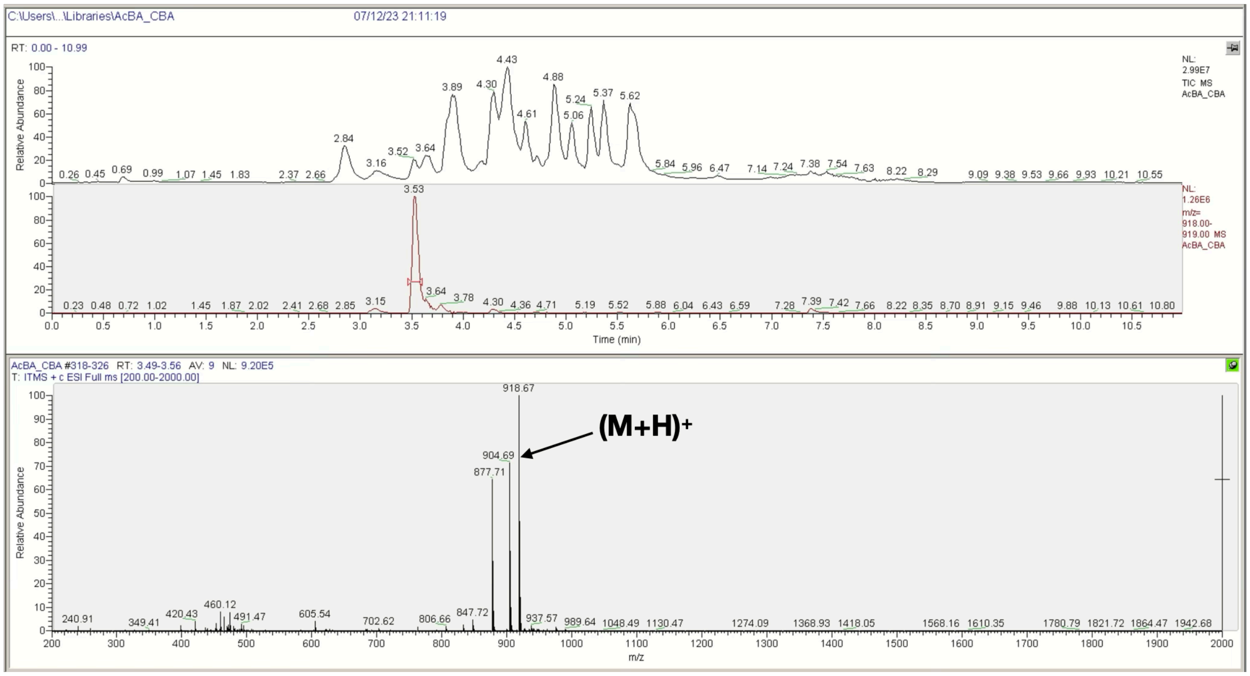


**F** **Ac-βA-RGEFV-Glu-βA-NH2**

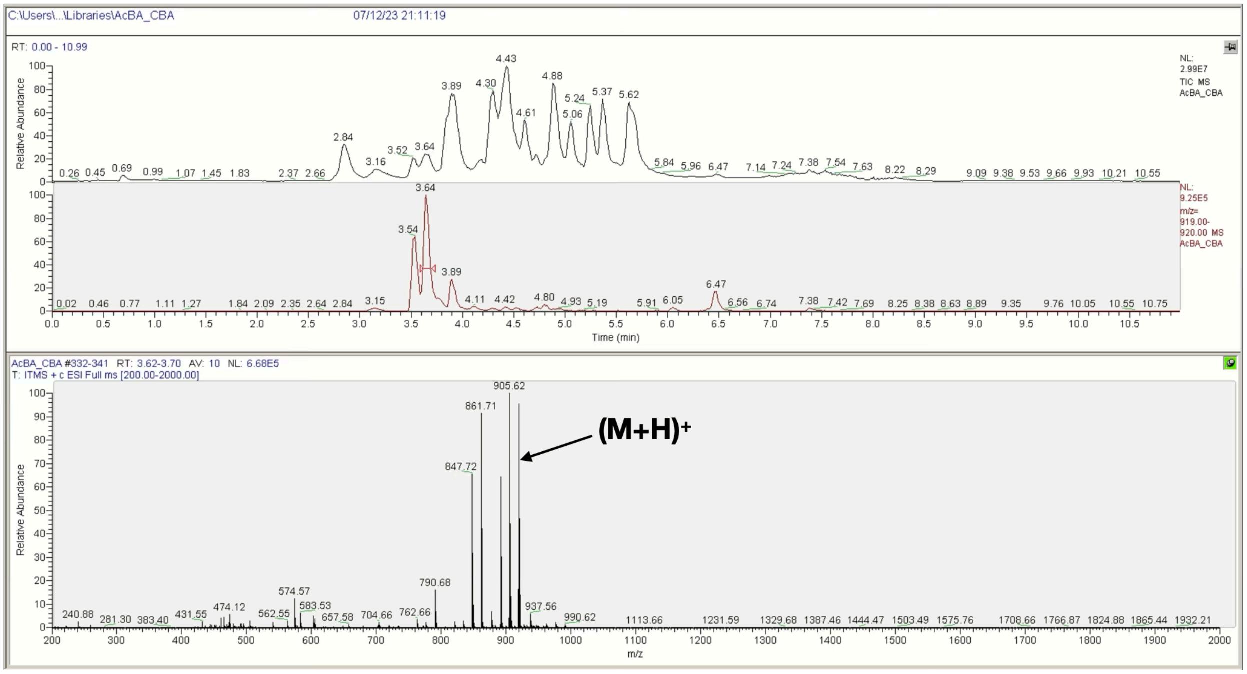

**G** **Ac-βA-RGEFV-Gly-βA-NH$_2$**

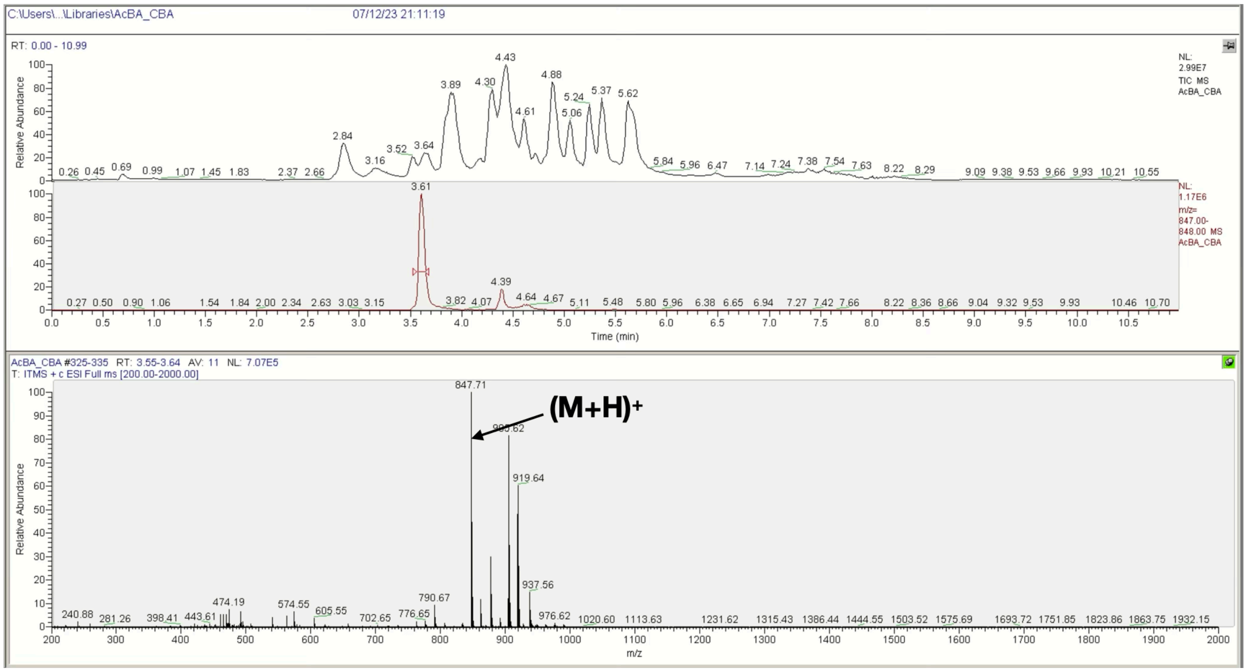


**H** **Ac-βA-RGEFV-His-βA-NH$_2$**

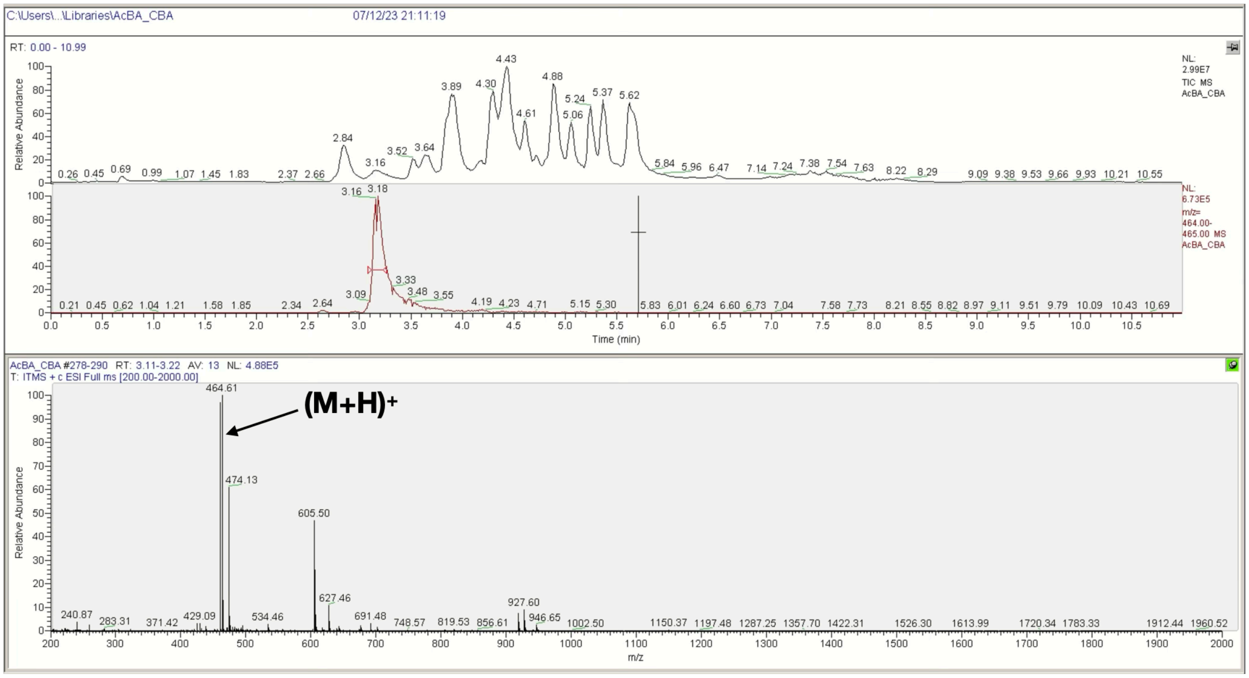

**I** **Ac-βA-RGEFV-Ile/Leu-βA-NH$_2$**

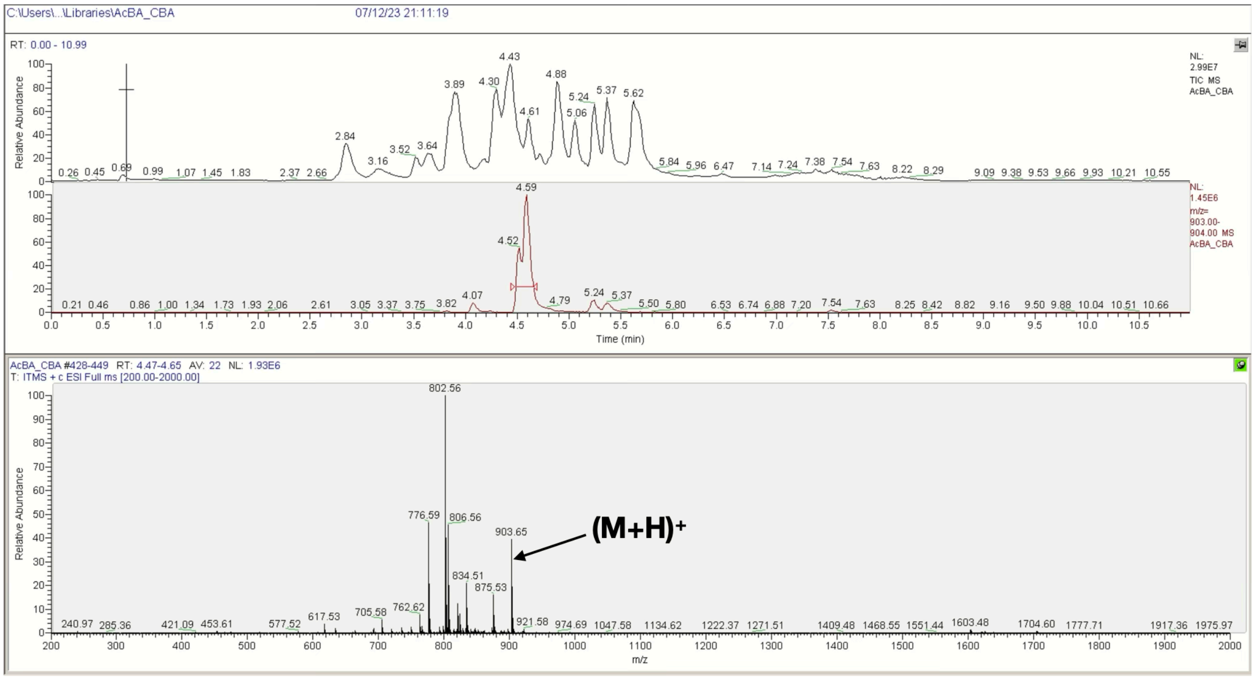


**J** **Ac-βA-RGEFV-Lys-βA-NH$_2$**

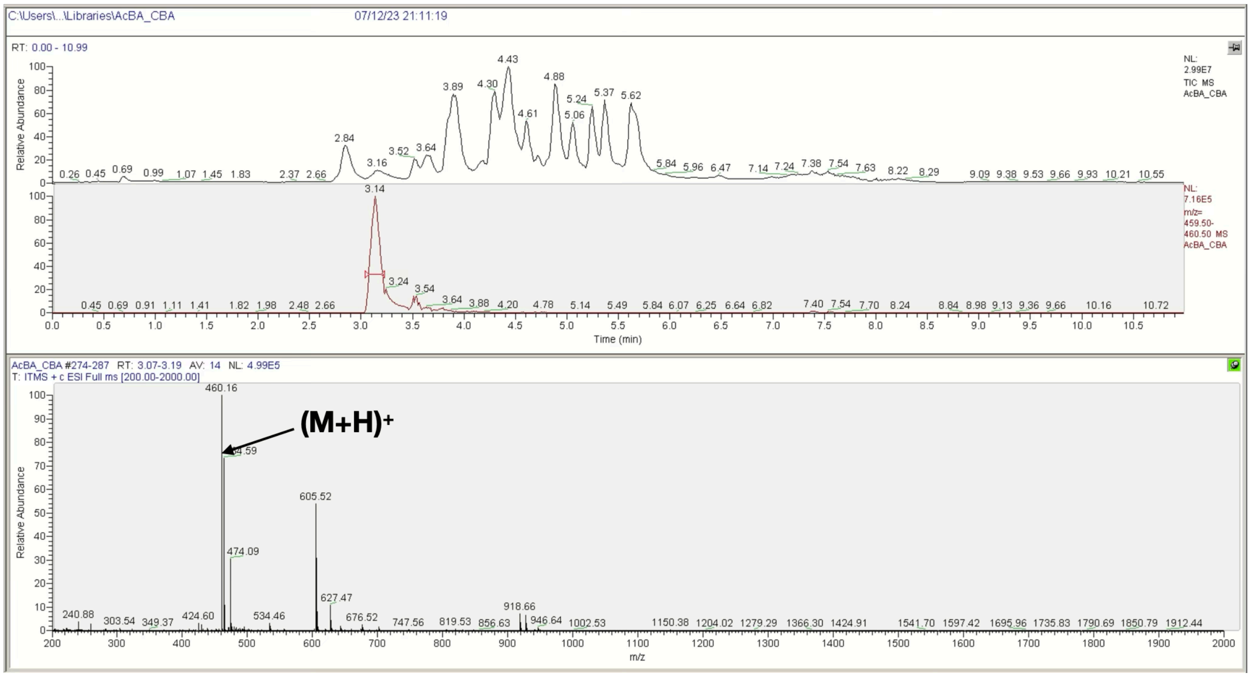

**K** **Ac-βA-RGEFV-Met-βA-NH2**

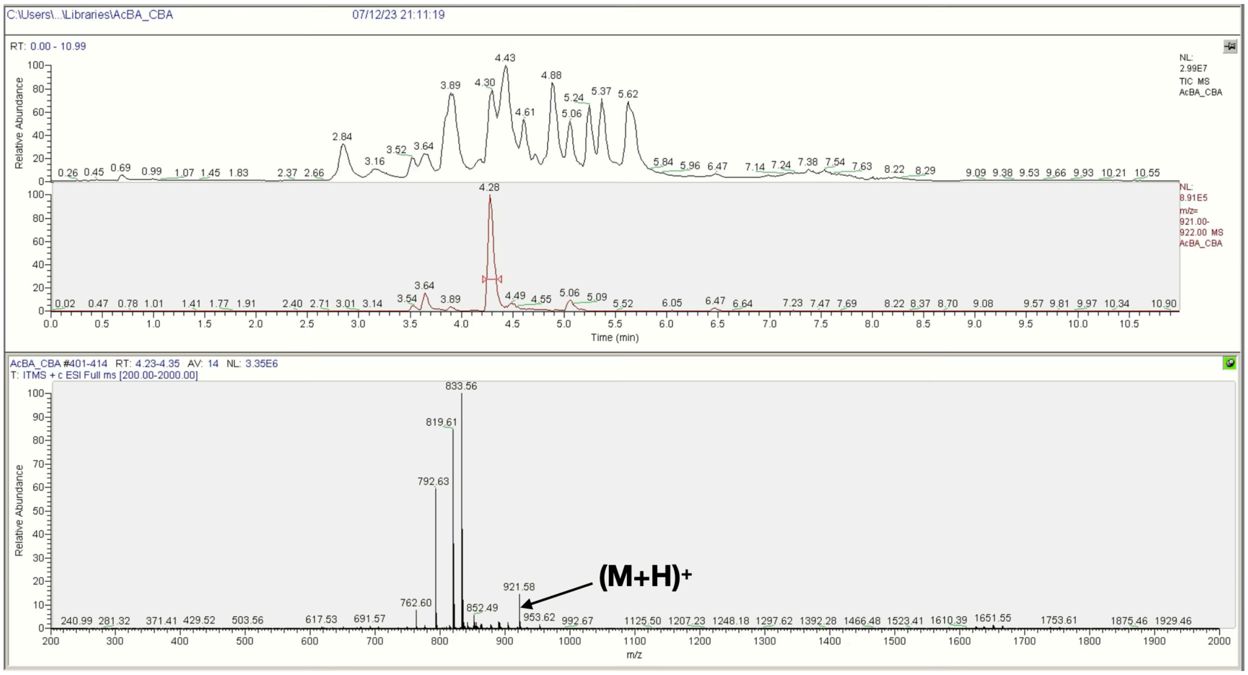


**L** **Ac-βA-RGEFV-Phe-βA-NH2**

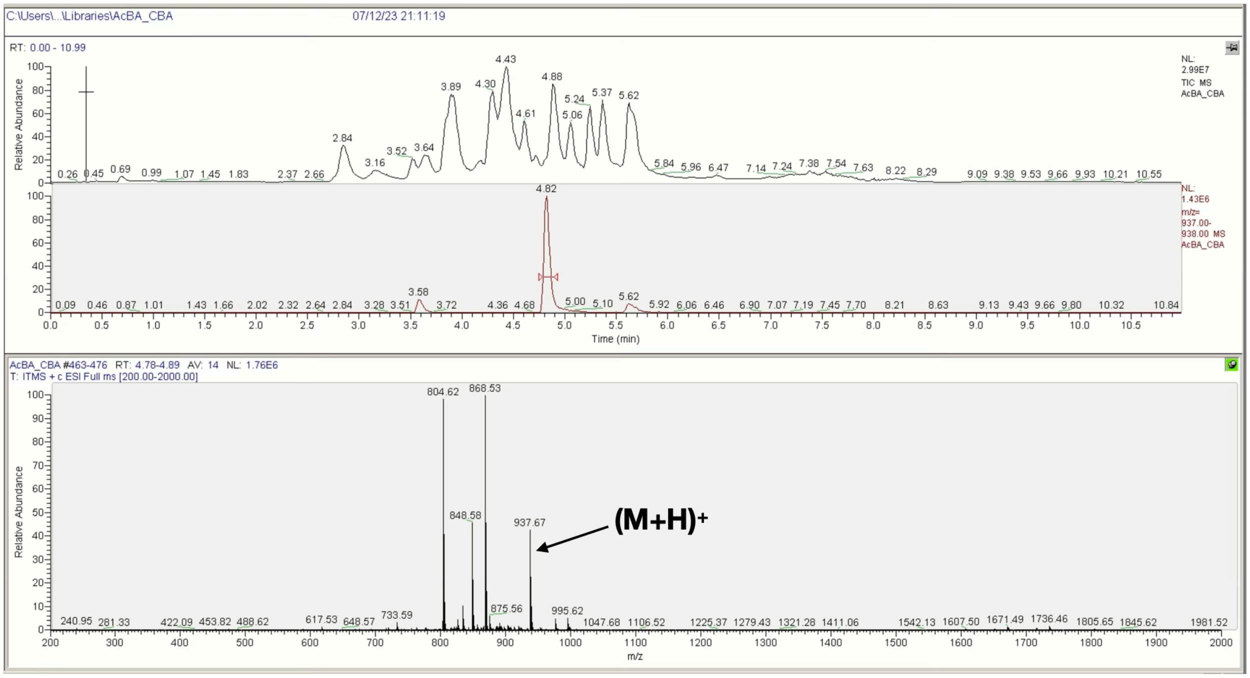

**M** **Ac-βA-RGEFV-Pro-βA-$NH_2$**

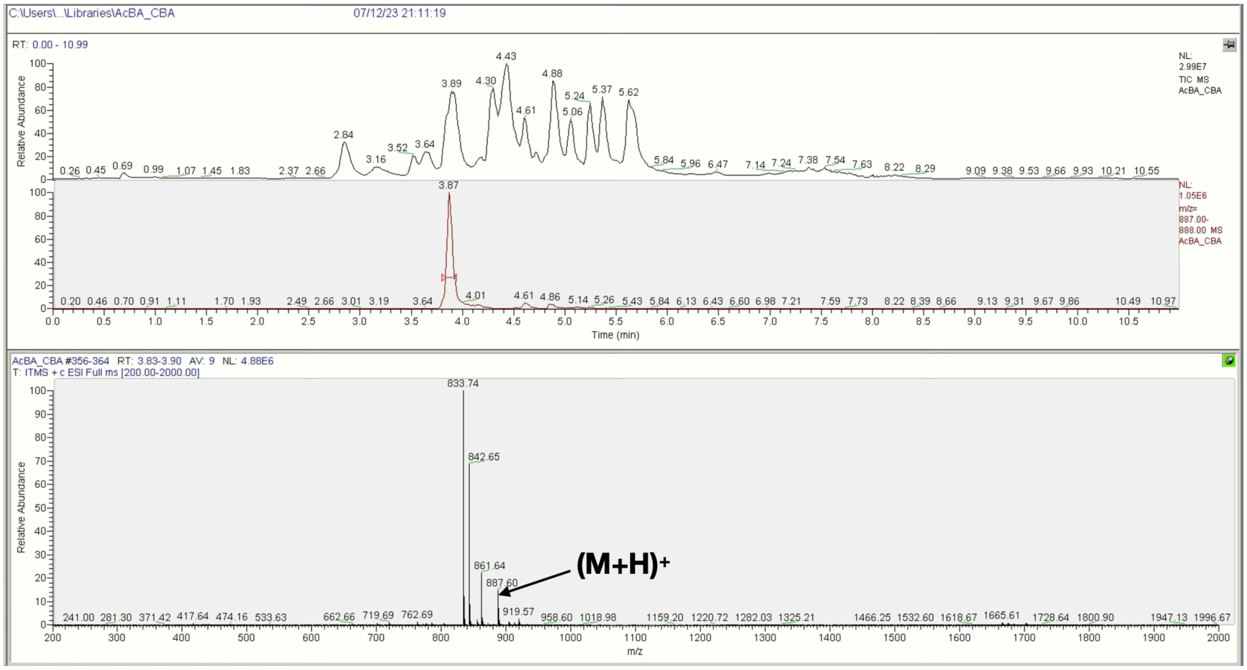


**N** **Ac-βA-RGEFV-Ser-βA-$NH_2$**

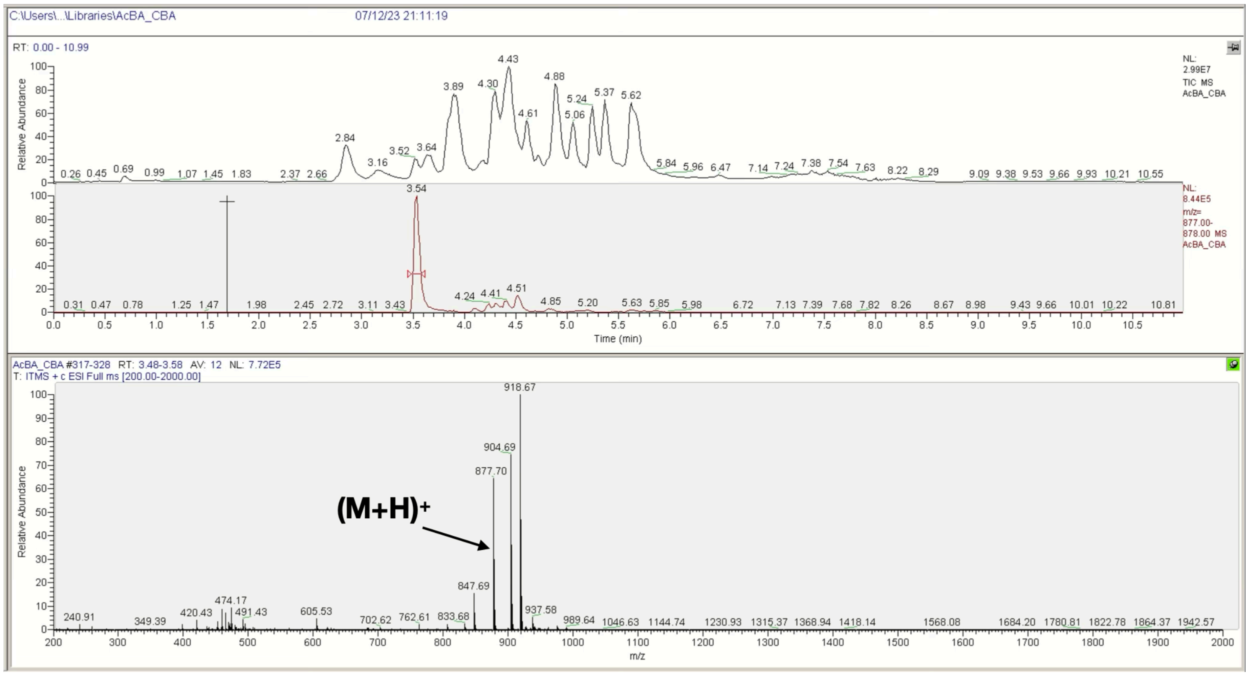

**O** **Ac-βA-RGEFV-Thr-βA-NH2**

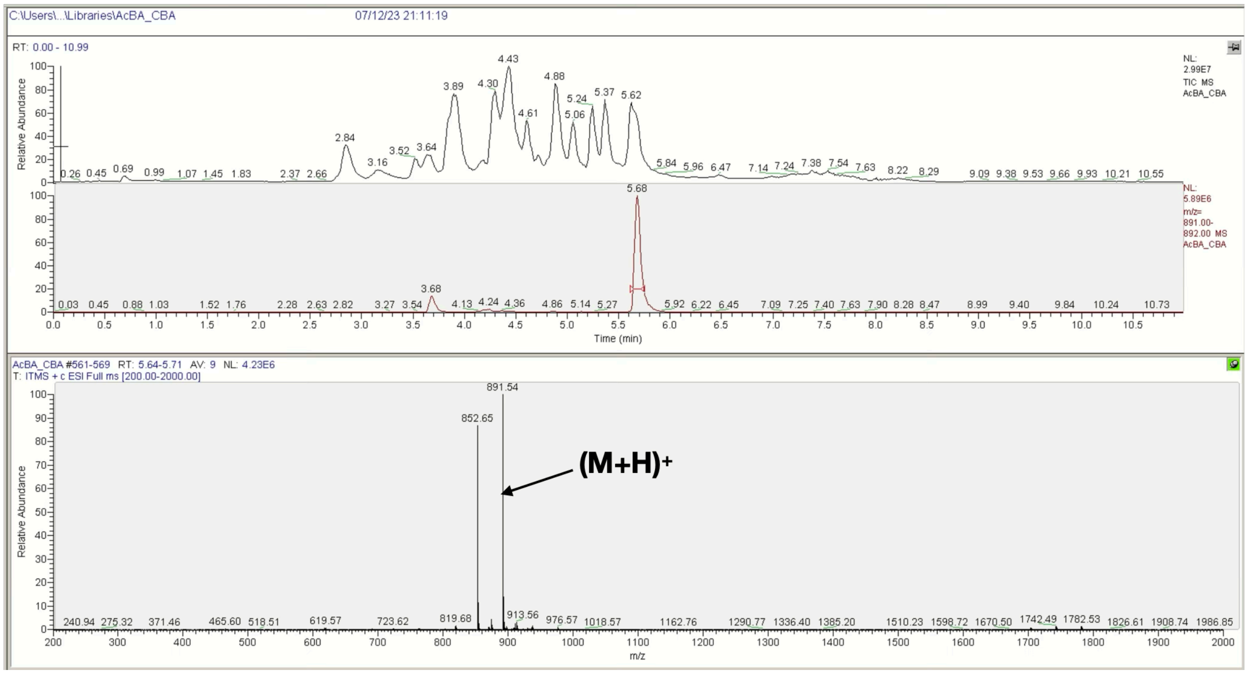


**P** **Ac-βA-RGEFV-Trp-βA-NH2**

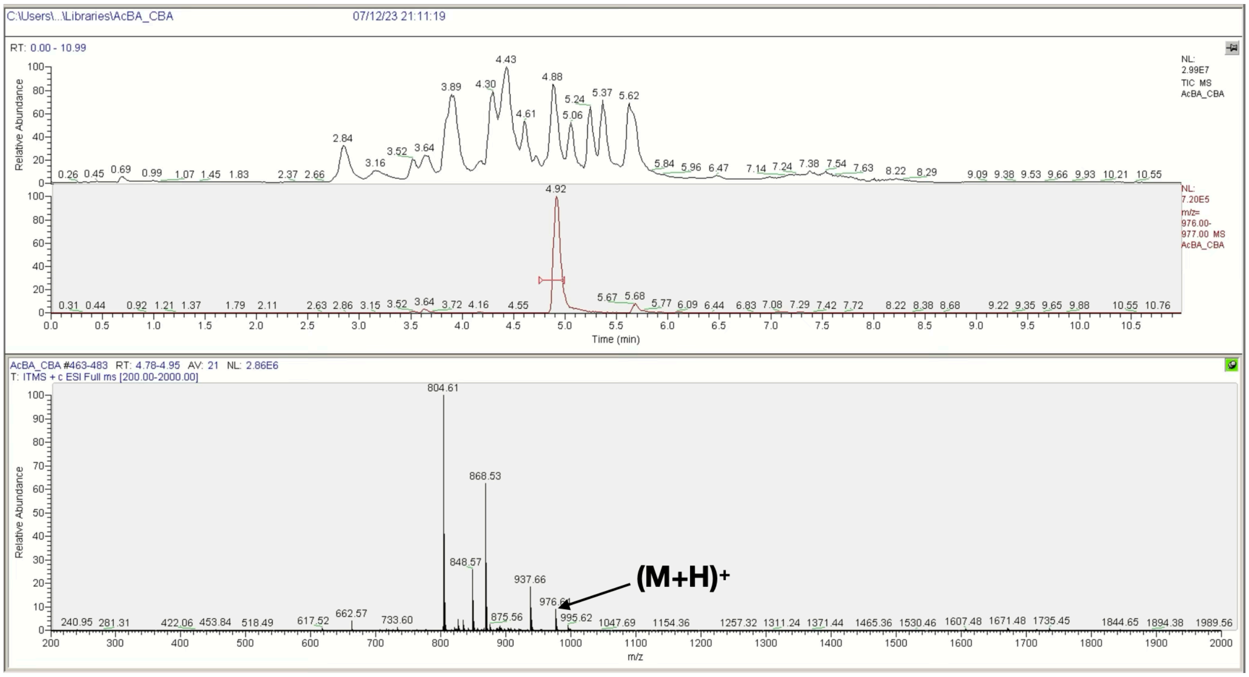

**Q** **Ac-βA-RGEFV-Tyr-βA-$NH_2$**

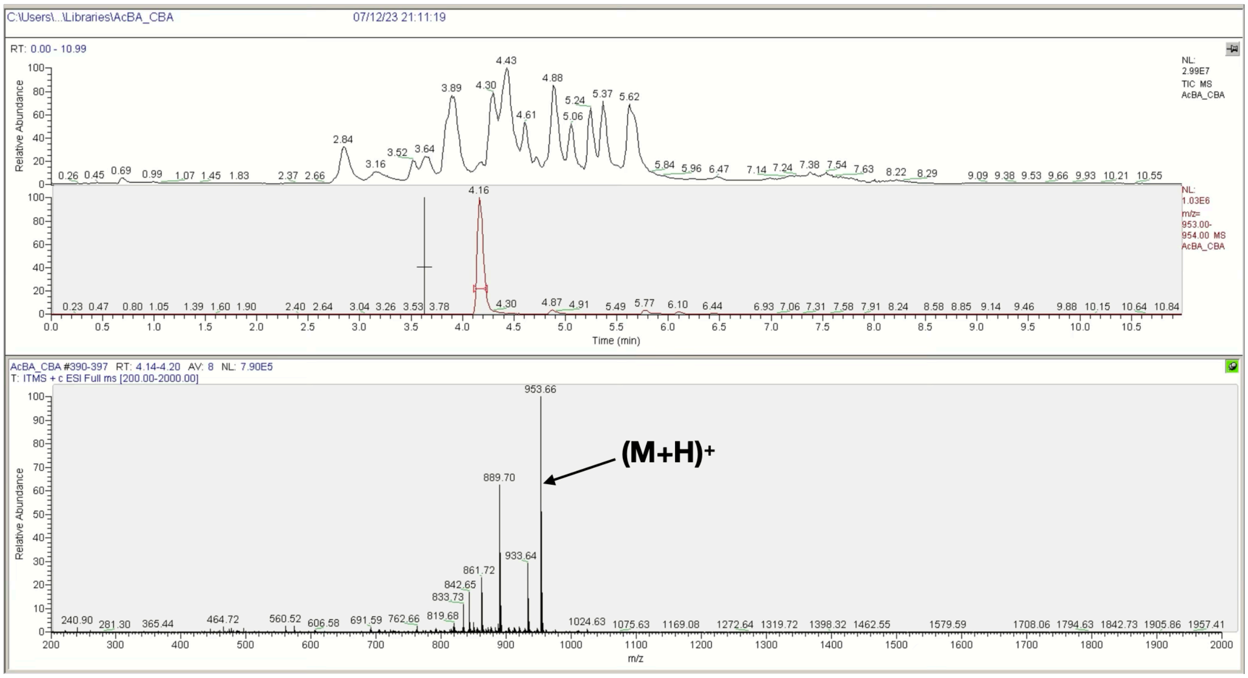


**R** **Ac-βA-RGEFV-Val-βA-$NH_2$**

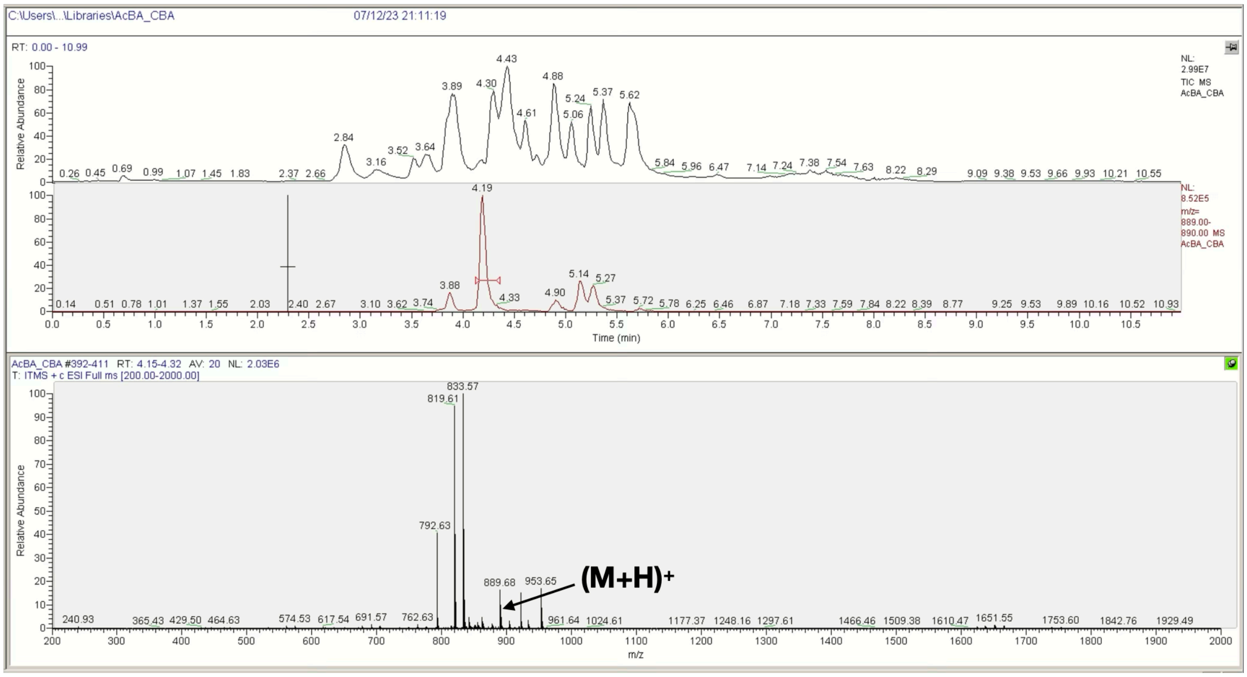


**Figure S3**. LCMS spectra of the Ac-βA-RGEFV-X-$NH_2$ libraries, where X = a) Ala, b) Arg, c) Asn, d) Asp, e) Gln, f) Glu, g) Gly, h) His, i) Ile/Leu, j) Lys, k) Met, l) Phe, m) Pro, n) Ser, o) Thr, p) Trp, q) Tyr, r) Val.

**$NH_2$-βF-(βA)$_6$-Amide**

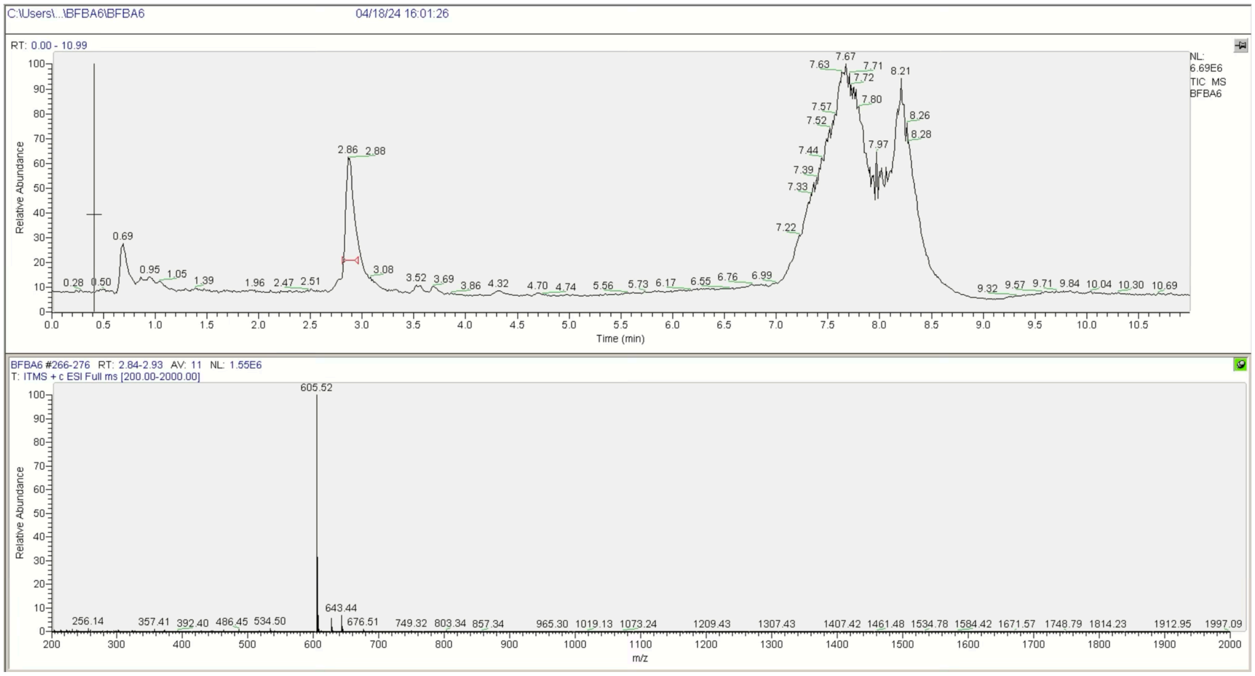


**Figure S4**. LCMS spectra of the internal standard $NH_2$-(βF)-(βA)$_6$-amide.
